# Single-cell characterization of the human immune landscape reveals dysfunction in monocyte-mediated cellular interactions in symptomatic malaria

**DOI:** 10.64898/2026.10.14.759097

**Authors:** Loréna Quatreville, Mathieu Adjemout, Charlyne Gard, Magali Torres, Camille Cohen, Brigitte Tumamo Fotso, Sandie Ménard, Abdel Yaya-Oye Akibou, Valérie Duplan-Eche, Anne-Laure Iscache, Martine Biarnes-Pelicot, Balotin Fogang, Antoine Berry, Nicolas Blanchard, Lionel Spinelli, Sandrine Nsango, Antoine Claessens, Sandrine Marquet

## Abstract

*Plasmodium falciparum* infection results in a wide range of phenotypes including chronic asymptomatic and symptomatic individuals. The malaria outcome depends, at least in part, on host immune responses mediated by specific cell subsets. Here, we investigated the immune landscape of chronic asymptomatic and symptomatic children using single-cell RNA sequencing. Unexpectedly, we uncovered biologically relevant immune cell complexes, which were further confirmed by imaging flow cytometry. Symptomatic infection was associated with a decrease of cell-cell interactions, particularly between non-classical monocytes and T cells. Non-classical monocytes from symptomatic children exhibited profiles with increased inflammation characterized by enhanced *S100A8*-*S100A9* expression together with decreased expression of interaction-associated genes such as *CD86* or *ITGAL*. Moreover, we demonstrated that inflammatory and cell interaction programs could not be activated simultaneously within the same cell. In addition, the concomitant expression of inflammatory genes and members of the *LILR* gene family was predominantly observed in monocytes from symptomatic individuals. These findings raise the possibility that enhanced *S100A8*/*S100A9*-*LILRB1* signaling may contribute to the altered functional state of monocytes in symptomatic malaria and, potentially, may impair their interactions with T cells. Together, our results suggest that immune cell complexes may therefore reveal an additional layer of immune regulation associated with the development of clinical malaria.

## Introduction

Malaria remains a major public health problem in endemic regions, causing substantial morbidity and mortality, particularly in sub-Saharan Africa. In 2024, an estimated 282 million malaria cases were reported worldwide, resulting in approximately 610,000 deaths, most of which occurred in children under five years of age ^1^. Despite widespread exposure in endemic areas, the clinical outcome of *Plasmodium falciparum* infection varies considerably among individuals. Infections may be naturally cleared, remain asymptomatic, or progress to clinical forms ranging from uncomplicated malaria to life-threatening severe malaria ^2^. This striking heterogeneity results from a complex interplay between host and parasite genetics, acquired immunity, host–pathogen interactions, and environmental characteristics. However, the mechanisms governing these diverse clinical trajectories remain incompletely understood, limiting the development of effective therapeutic and preventive strategies.

*P. falciparum* infection induces a complex and dynamic immune response involving both innate and adaptive compartments. Although effective immune activation is necessary for parasite control, excessive or dysregulated inflammation contributes to disease severity. In malaria-endemic regions, repeated exposure, especially during childhood, gradually leads to the acquisition of partial clinical immunity as reflected by an increased proportion of asymptomatic individuals with age ^3–5^. Long-lasting asymptomatic infections are thought to rely on disease tolerance mechanisms that limit immunopathology despite persistent parasitemia. These individuals also exhibit antibodies against RBCs, high levels of IgG, and almost no signs of inflammation ^6^. They also display durable changes in the frequency and immunoregulation of circulating innate and adaptive T lymphocytes. Several studies have identified immune features associated with asymptomatic *P. falciparum* infection, including increased frequencies of PD-1+ exhausted CD4+ T cells, activated regulatory T cells, and IL-10 responses, supporting an immunoregulatory profile ^7,8^. Repeated malaria exposure has also been associated with the progressive loss and functional impairment of Vδ2+ γδ T cells, which may contribute to immune tolerance and reduced inflammatory pathology ^9^. In the B cell compartment, atypical memory B cells expand during malaria, although their functional significance remains controversial ^10–13^. Despite these advances, the molecular programs underlying these immune adaptations remain largely unresolved.

Bulk transcriptomic studies have shown that symptomatic *P. falciparum* infection induces changes in host gene expression, characterized by activation of innate immune, interferon signaling, cytokine production and inflammatory pathways, together with transcriptional signatures related to erythropoiesis, metabolic stress, and immune regulation ^14^. These transcriptional profiles reflect both parasite-driven pathology and host adaptive responses to infection. Severe malaria is associated with exacerbated inflammatory responses and dysregulation of endothelial and coagulation pathways, whereas mild disease is characterized by a more balanced transcriptional program combining immune activation with regulatory mechanisms. In contrast, asymptomatic *P. falciparum* infection has been reported to elicit no detectable host transcriptional response in peripheral blood, suggesting that the parasite may remain below the host’s clinical and immunological radar ^15,16^. However, these results are based primarily on bulk transcriptomic analyses, which average gene expressions across heterogeneous cell populations. Consequently, they obscure cell type-specific responses and fail to capture rare but potentially critical immune cell subsets. As a result, the cellular and molecular mechanisms that determine whether infection leads to protection, asymptomatic carriage, or clinical disease remain poorly defined. Single-cell RNA sequencing (scRNA-seq) now provides the opportunity to resolve this complexity by profiling transcriptional programs at cellular resolution and identifying immune populations associated with protection or immunopathology.

To date, only a limited number of studies have applied scRNA-seq to investigate host immune responses during malaria infection ^17–20^. These studies have revealed extensive remodeling of the immune landscape, including the expansion of immunoregulatory monocytes, IL-10–producing Tr1 CD4 T cells, regulatory B cells, and NK and γδ T cells with tolerogenic features ^18^, together with a widespread type I interferon signature across multiple immune cell subsets. They have also uncovered inter-ethnic differences in immune responses, demonstrating that naturally malaria-resistant populations display distinct cell type-specific transcriptional programs ^17^.

Here, we performed single-cell RNA sequencing on peripheral blood mononuclear cells (PBMCs) from children enrolled in a longitudinal cohort study who were infected with *P. falciparum* and presented with either persistent asymptomatic infection (chronic individual) or symptomatic malaria. We identified for the first time in malaria, unexpected subsets of interacting cells corresponding to monocyte-T cell complexes that were significantly more abundant in children with chronic infection and characterized distinct immune signatures that distinguish the two clinical groups. Analysis of these subsets revealed impaired interactions between non-classical monocytes and T cells in symptomatic children, which were associated with an exacerbated inflammatory response that may prevent T cell activation. In addition, a concomitant inflammatory response characterized by *S100A8* and *S100A9* expression, together with expression of gene from the *LILR* family, may contribute to malaria pathogenesis.

## Results

### Study overview and patient population

To characterize the immune responses associated with symptomatic and chronic asymptomatic malaria infection, we combined droplet-based single-cell RNA sequencing (scRNA-seq), and flow cytometry. High-quality peripheral blood mononuclear cell (PBMC) samples from three children with symptomatic malaria and four with chronic asymptomatic infection were selected for single-cell transcriptomic analysis (Fig. 1a and Supplementary Table 1). Venous blood samples from symptomatic individuals were collected at symptom onset, whereas samples from individuals with chronic asymptomatic infection were collected 3–4 weeks after the first *P. falciparum*-positive detection. Chronic asymptomatic infection was defined as persistent *P. falciparum* infection lasting more than 22 days according to the classification recently established by Fogang *et al.* (manuscript in preparation). In order to preserve PBMC transcriptomic profiles integrity, we used a probe-based approach for fixed samples. Following initial quality control, a total of 69,533 high-quality cells were retained for subsequent single-cell transcriptomic analyses.

**Figure 1.**
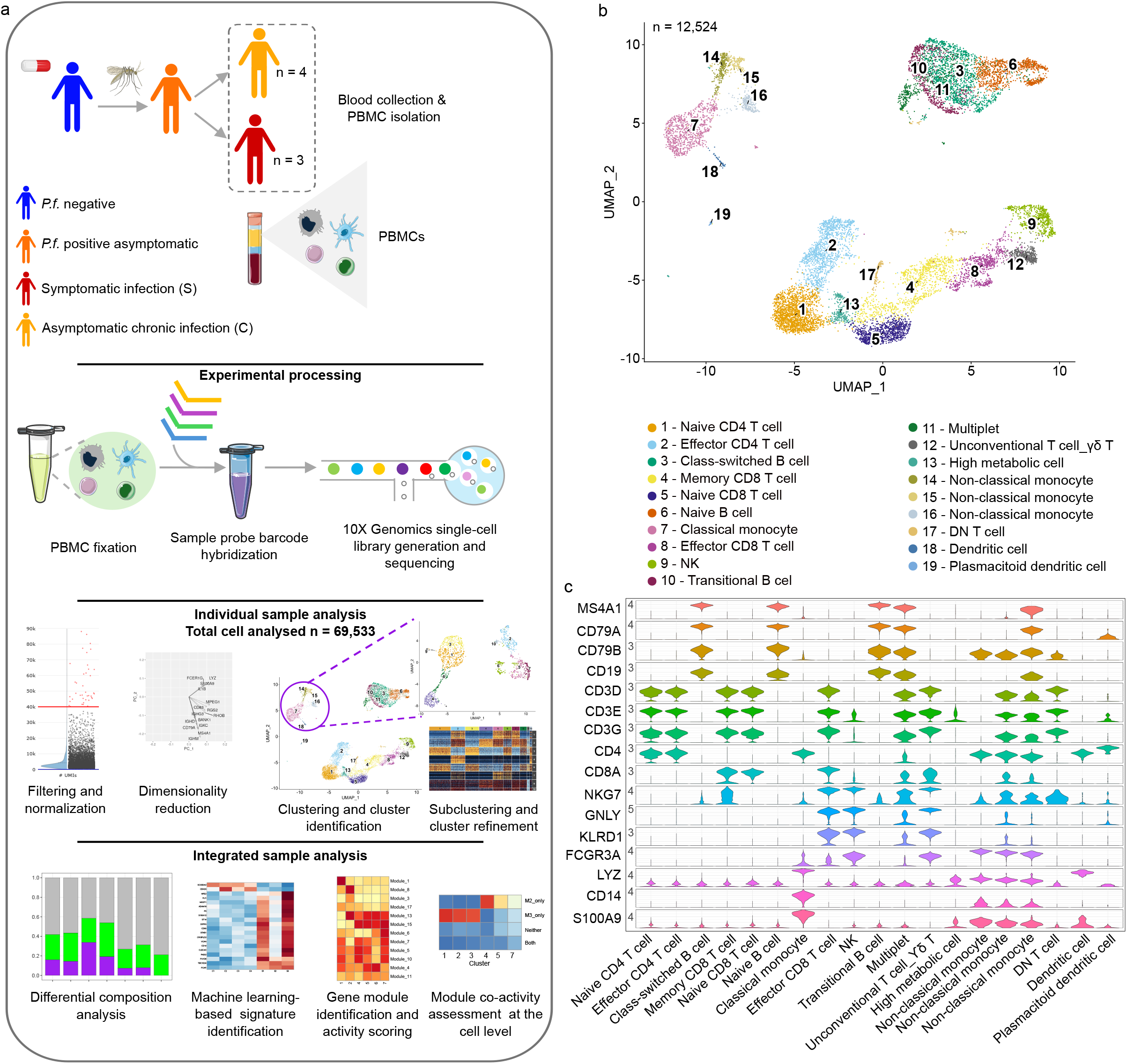
Experimental design and identification of major cell populations by single-cell RNA sequencing. a Schematic overview of the experimental design and analytical workflow, including sample processing, single-cell RNA sequencing, and bioinformatic analyses. b-c Representative UMAP projection (b) and corresponding marker-gene expression violin plots (c) used to identify major cell populations in a C2 representative individual. UMAP was generated after the clustering and marker gene identification (FindAllMarkers) steps from the Seurat standardized pipeline. The violin plot shows the abundance of cells expressing specific marker genes of B cells, T cells and monocytes selected for visualization.

### Circulating monocytes are increased in symptomatic malaria

Unsupervised clustering of the scRNA-seq data identified between 14 and 20 distinct cell clusters per individual (Supplementary Fig. 1). Cell identities were assigned based on publicly available reference databases, including PanglaoDB, and previously published immune cell signatures ^21,22^. Cluster annotation was visualized using UMAP (Uniform Manifold Approximation and Projection), with the representative sample C2, comprising 19 clusters, shown in Fig. 1b. Overall, the dataset included five myeloid cell clusters (monocytes: clusters 7, 14, 15, and 16; dendritic cells, cluster 18), three B cell clusters (clusters 3, 6, and 10), seven T cell clusters (clusters 1, 2, 4, 5, 8, and 12) and natural killer (NK) cell cluster (cluster 9) (Fig. 1b and c). We also identified a cluster enriched for metabolically active cells (cluster 13) and two rare cells populations corresponding to double-negative T cells (cluster 17) and plasmacytoid dendritic cells (cluster 19).

To determine whether differences in immune cell composition were associated with clinical outcomes, we compared the relative abundance of each immune cell population between symptomatic and chronic asymptomatic malaria. There was no significant difference in the proportions of effector-memory CD8^+^ T and NK cells, effector-memory CD4^+^ T cells, and B cell clusters between the two groups (Fig. 2a-c). Although the difference did not reach statistical significance, symptomatic children tended to exhibit lower frequencies of CD8⁺ T cells than children with chronic infection (Fig. 2a). In contrast, symptomatic children showed a significantly higher proportion of circulating monocytes (P = 1.3 x 10^-3^; Fig. 2d and e) compared with chronic asymptomatic children. This increase was independently confirmed by flow cytometry on the same samples (P < 1 x 10^-3^; Fig. 2f), showing concordance between the two experimental approaches. Together, these findings identified circulating monocytes as the immune cell population most strongly associated with symptomatic malaria, suggesting that monocyte abundance is associated with clinical outcome.

**Figure 2.**
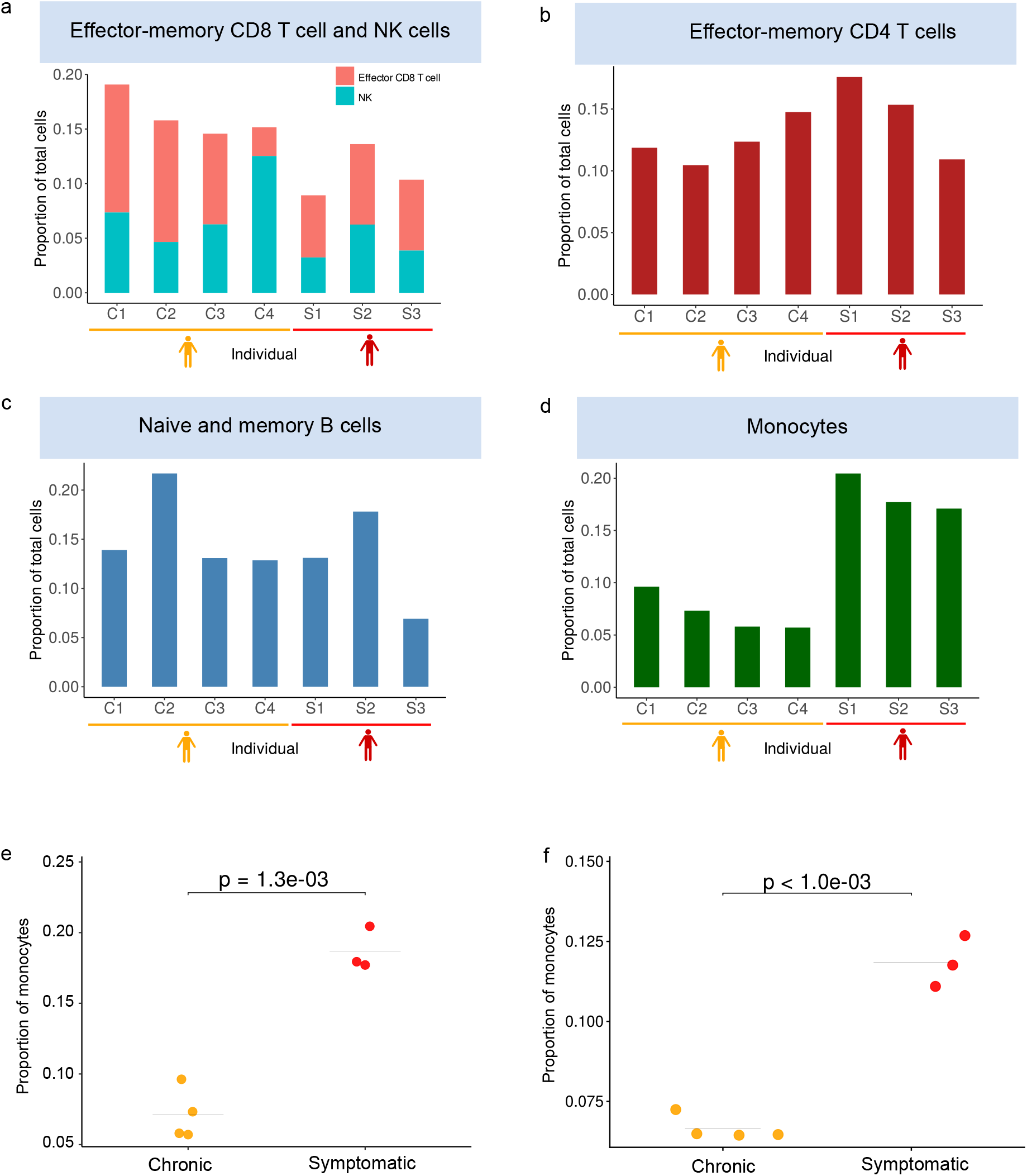
Identification of major and rare cell populations in chronic and symptomatic individuals. a–d Proportions of major immune cell populations across all individuals, including effector and memory CD8 T and NK cells(a), effector CD4 T cells (b), naive-memory B cells (c), and monocytes (d). e-f Proportion of monocytes in chronic and symptomatic individuals from single-cell RNA-Seq data (e) and flow-cytometry (f). Statistical significance was assessed with a beta-binomial model using DCATS (*FDR* = 1.3 x 10⁻^3^) and sccomp (*FDR* < 10^-3^) respectively.

### Subclustering analyses did not identify cell population subtypes specific to symptomatic malaria

To further dissect the cellular heterogeneity within each major immune lineage, we performed for each individual separately, subclustering analyses based on co-expression gene modules using SciGeneX algorithm (Fig. 3a, Supplementary Table 2). Such analyses were performed for all children, and the corresponding results are presented in Supplementary Table 3. As illustrated for individual C2, we identified a total of 1169 informative genes for effector-memory CD8+ T and NK cells, distributed across 11 distinct modules. Briefly, module 1 was predominantly expressed in clusters 5, 6, 7 and 8, while module 3 showed preferential expression in cluster 2 and 11. In contrast, module 2 exhibited higher expressions in clusters 1, 2, 3, 9 and 10 (Fig. 3a). We can visualize on the UMAP individual module expression using an averaging module score (Fig. 3b). As illustration, cells exhibiting high module 2 activity predominantly corresponded to clusters 1, 2, and 3, while cells with high module 3 activity were mainly associated with clusters 2 and 11. Importantly, evaluating combinations of module activities allowed us to identify and discriminate transcriptionally distinct cell populations that may otherwise be grouped within the same cluster (Fig. 3c). For example, cluster 2 exhibits both a cytotoxic signature and a CD56 dim NK signature allowing to identify this cluster as a CD56 dim NK population with cytotoxic properties. This module-based approach therefore provides a higher-resolution characterization of cellular heterogeneity and enables a more refined delineation of effector-memory CD8^+^ T and NK cells, effector-memory CD4^+^ T cells, naïve-memory B cells, and monocyte subpopulations (Fig 3a-f). In the representative sample C2, the CD8+ T/NK-cell compartment identified eleven clusters, compared with only four clusters (clusters 4, 8, 9, and 12) identified in the initial analysis. This higher resolution allowed the identification of three NK cells subsets (CD56 dim NK1-cytotoxic, CD56 bright NK2 and CD56 dim NK3-cytotoxic) as well as rare lymphocyte populations such as γδ T cells and NKT cells (Fig. 3a and c). Likewise, the effector memory CD4+ T-cell compartment previously represented by a single CD4+ T-cell cluster (cluster 2), was resolved into six transcriptionally distinct populations corresponding to major functional subsets, including Th1, Th2, Th17 and regulatory T (Treg) cells (Fig. 3d). Similarly, the B-cell compartment expanded from four clusters (clusters 3, 6, 10, and 11) to seven distinct populations (Fig. 3e). The monocyte compartment exhibited an even greater increase in complexity, expanding from four clusters (clusters 7, 14, 15, and 16) to ten clusters corresponding to transcriptionally distinct populations comprising multiple classical and non-classical monocyte subsets, as well as a distinct intermediate monocyte population (Fig. 3f). This strategy provided increased cellular resolution across all seven individuals (Supplementary Table 4). High-resolution subclustering analyses did not reveal any immune cell subtype that was consistently associated with either clinical phenotype, including among rare cell populations captured through the comprehensive cellular profiling of our dataset (∼10,000 cells per individual). Instead, both immune cell composition and transcriptional states showed marked inter-individual variability, highlighting the biological heterogeneity and complexity of the host’s immune response to *Plasmodium falciparum* infection.

**Figure 3.**
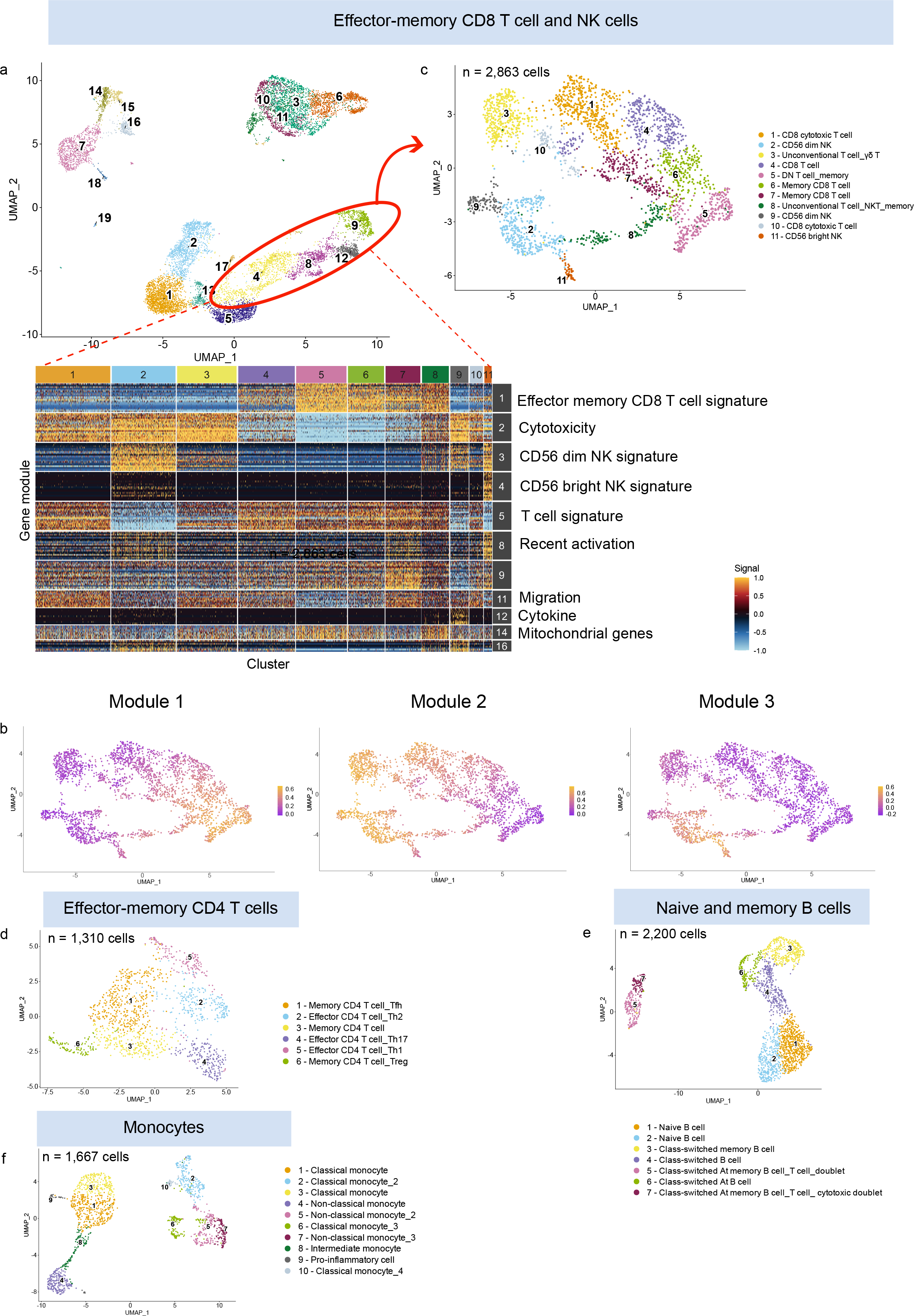
Identification of rare cell types through cluster refinement in chronic and symptomatic individuals. a UMAP projection showing effector CD8 T and NK cells subclustering and heatmap displaying the modules of co-expressed genes used for subcluster annotation. b UMAP projection or module scores corresponding to module 1, module 2 and module 3. c UMAP projection of the subclusters identified with the heatmap. d-f UMAP projections showing cell populations identified following subclustering and module gene identification for effector-memory CD4 T cells (d), naive-memory B cells (e), and monocytes (f). Results are shown for one representative individual (C2).

### Unexpected identification of interacting cells as subclusters of monocytes

Unexpectedly, monocytes segregated into two major transcriptional clusters that were independent of the canonical classification into classical and non-classical monocytes, with both subsets distributed across the two clusters (Fig. 3f). This observation suggested that monocyte diversity extends beyond the conventional classification and prompted us to further investigate the transcriptional programs defining these cellular states. A more detailed examination of the gene list modules within the monocyte compartment uncovered unexpected subpopulations co-expressing transcriptional programs belonging to different immune lineages (Fig. 4a, Supplementary Table 2). Specifically, clusters 5, 6, and 7 co-expressed canonical monocyte and T cell gene modules, whereas a small subset of cells in cluster 10 co-expressed monocyte and B cell genes. Strikingly, cells in cluster 2 exhibited the broadest transcriptional repertoire, simultaneously expressing gene modules characteristic of monocytes, T cells, and B cells, revealing an unexpected level of transcriptional complexity within the monocyte compartment. In addition, Module 6 was enriched for genes associated with activated cellular states, a functional signature shared by clusters 1, 3, 6, 8, 9 and 10. A distinct cytotoxicity-related signature was observed across clusters 2, 5, and 6. Accounting for these complex subsets enabled a more accurate annotation of monocyte subclusters (Fig. 4a). Such co-expression profiles observed in the hybrid transcriptome clusters (2, 5, 6, 7 and 10) strongly indicate the presence of multiplets, including doublets or triplets resulting from the co-encapsulation of multiple cells within a single droplet. Looking at all cell types, we identified a proportion of multiplets ranging from 6.2 % to 15.5 % (Supplementary Table 5), exceeding the multiplet rate predicted by the 10x Genomics platform (approximately 8% for 10,000 recovered cells). Importantly, the rate reported by 10x Genomics includes both embedded multiplets, composed of cells from the same lineage, and neotypic multiplets, composed of distinct cell types ^23^. Since only multiplets involving different cell types could be reliably identified and quantified in our samples, the frequency of doublets and triplets in our dataset is likely underestimated. To account for this limitation, we estimated the number of embedded multiplets mathematically. When these estimated embedded multiplets were combined with the experimentally identified neotypic multiplets, the overall multiplet frequency substantially exceeded the rate expected from the 10x Genomics model, reaching up to 17% (Supplementary Table 6). Although a proportion of these events is expected to represent technical multiplets generated by the random co-encapsulation of two or more independent cells, the marked excess over the predicted rate suggests that a subset of the detected multiplets likely corresponds to biologically associated cell complexes. Notably, multiplets involving monocytes represented the majority of neotypic multiplets. Furthermore, their frequency was significantly higher than expected under a binomial model of random cell co-encapsulation (Supplementary Table 6), providing additional evidence that these associations are unlikely to result solely from stochastic encapsulation events. Indeed, these multiplets were independently validated by both optical bright-field and fluorescence microscopy (Supplementary Fig. 2). Monocyte-containing doublets and triplets were detected at multiple stages of sample processing, including both before and after PBMC fixation. Their presence prior to encapsulation demonstrates that these cell aggregates existed before droplet generation, supporting the conclusion that they represent genuine cell-cell complexes rather than technical artefacts generated during the encapsulation process.

**Figure 4.**
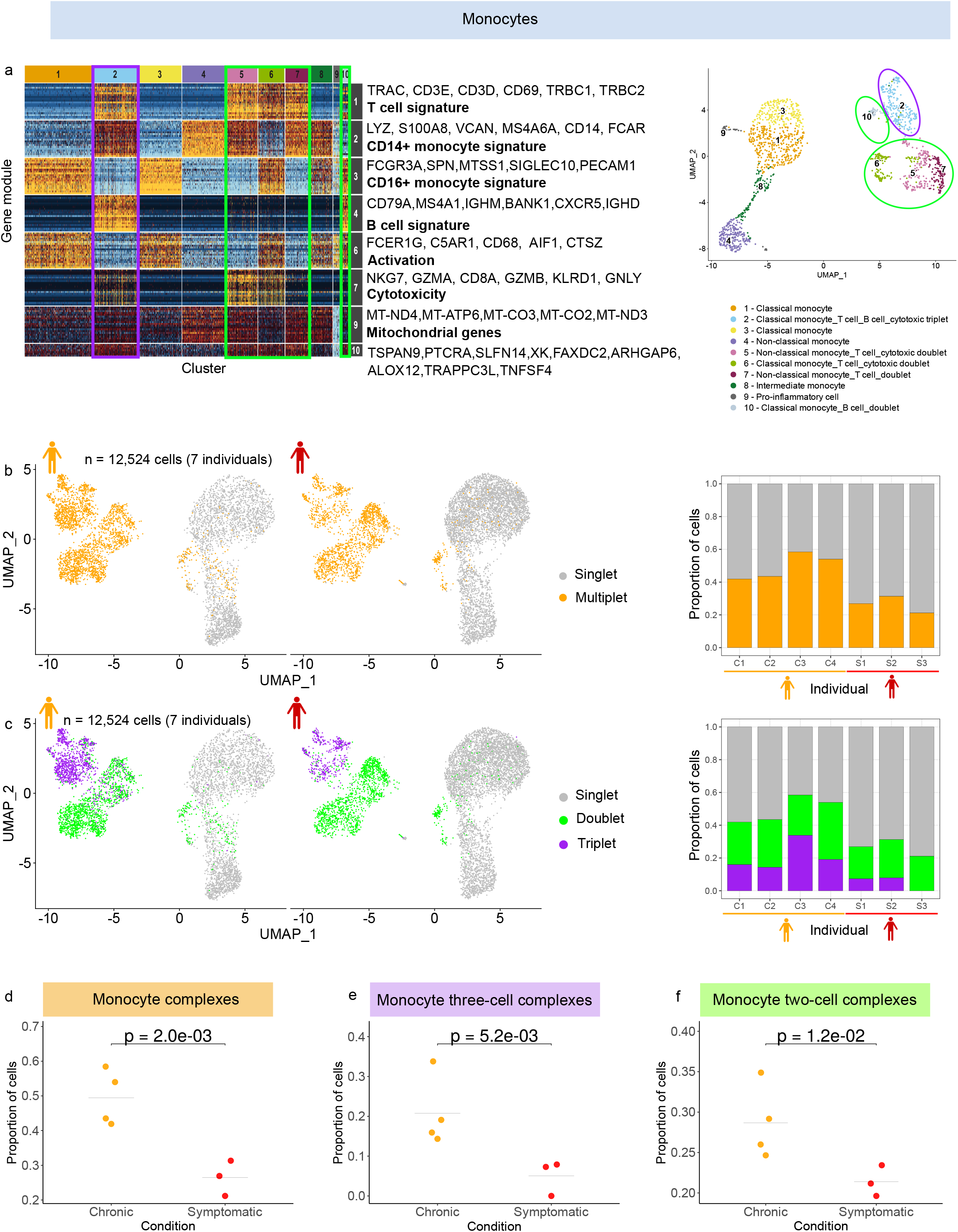
Monocyte complexes are significantly reduced in symptomatic individuals. a Heatmap of module genes and corresponding UMAP projection of monocyte populations identified. Major marker genes of each module are displayed on the right of heatmap.b UMAP projection and corresponding proportions of singlet and monocyte complexes across individuals. c UMAP projection and corresponding proportion of singlet, two-cell and three-cell monocyte complexes across individuals. d-f, Proportions of monocyte complexes (d), three-cell monocyte complexes (e) and two-cell monocyte complexes (f) in chronic asymptomatic and symptomatic individuals. Significant difference between conditions was assessed using DCATS with a beta-binomial model (*FDR* = 2.0 × 10⁻^3^, 5.2 × 10⁻^3^ and 1.2 × 10⁻^2^ respectively).

### Proportion of monocyte cell complexes distinguish symptomatic from chronic malaria

To investigate the biological relevance of these immune cell complexes, we evaluated whether their frequency differed between symptomatic and chronic malaria infection. The number of monocyte-containing complexes ranged from 403 to 857 per individual, corresponding to 3.9-8.2 % of the total cells analyzed, and comprised exclusively neotypic complexes (Fig. 4b and 4c). Complexes containing either three or two cells were on average 1.8-fold more frequent in asymptomatic children with chronic infection than in symptomatic children (P= 2.0 x 10^-3^, DCATS statistical test) (Fig. 4d). The proportion of three-cell complexes was significantly higher in asymptomatic children with chronic infection than in symptomatic children (P= 5.2 x 10-3) (Fig. 4e), as well as the proportion of two-cell complexes (P= 1.2 x 10^-2^) (Fig. 4f). Interestingly, symptomatic child S3, who had the highest fever, had no detectable three cell complexes.

Notably, despite monocyte increase in symptomatic individuals, the absolute number of monocyte complexes was similar between the two clinical groups, which may reflect a lower rate of complex formation in symptomatic children (Supplementary Table 5). We estimated that this rate was 2.7 times higher in chronic asymptomatic individuals.

We further investigated the cellular composition of these complexes to determine whether specific monocyte subsets were preferentially involved in cell-cell interactions and associated with clinical phenotype (Fig. 5a). Classical, intermediate and non-classical monocytes were detected within immune cell complexes. Symptomatic children had significantly lower proportions of complexes containing classical monocytes (P= 1.6 x 10^-2^, Fig. 5b) and non-classical monocytes (P=1.6 x 10^-2^, Fig. 5c) than children with chronic infection. Although complexes involving intermediate monocytes were more frequent in symptomatic children, this difference did not reach statistical significance (Fig. 5a). To identify the cell types that preferentially interacted with monocytes, we further characterized the composition of two and three cell complexes across individuals (Fig. 5d and 5e). Both classical and non-classical monocytes were involved in a broad range of complex configurations. Among these, complexes containing non-classical monocytes and T cell were significantly more abundant in children with chronic infection than in symptomatic children (P= 2 x 10^-2^) (Fig. 5d-f). Complexes composed of a non-classical monocyte, a T cell, and a B cell also tended to be more frequent in children with chronic infection (Fig. 5d and e). These complexes were predominantly composed of CD4^+^ T cells simultaneously interacting with a monocyte and a B cell.

**Figure 5.**
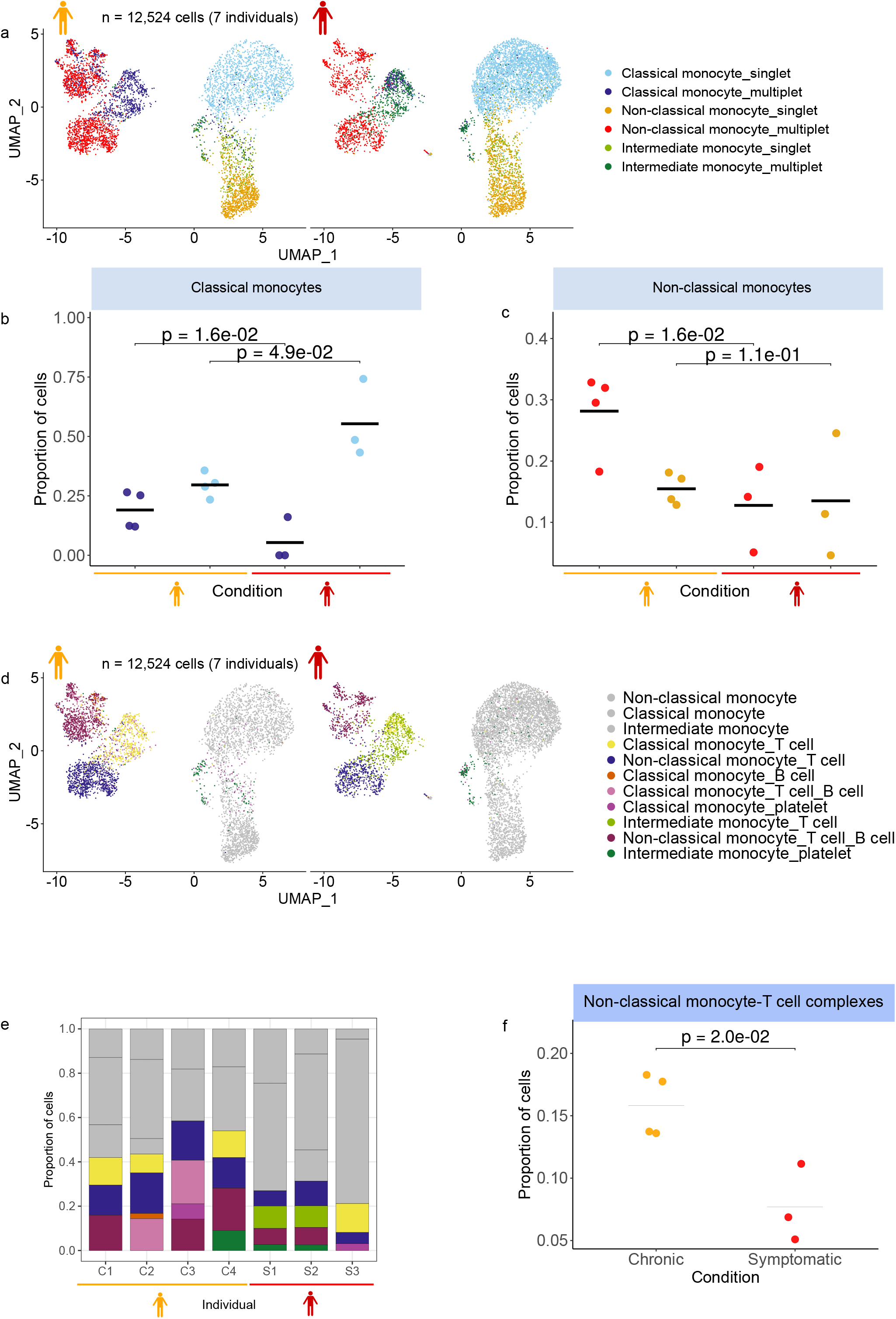
The proportion of monocyte subpopulation complexes is significantly lower in symptomatic individuals. a UMAP projection showing singlet and monocyte complexes stratified by classical, intermediate and non-classical monocyte identities in chronic asymptomatic and symptomatic individuals. b Proportions of classical monocyte singlets and complexes in chronic asymptomatic and symptomatic individuals. Statistical significance was assessed using DCATS with a beta-binomial model (*FDR* = 1.6 × 10^−2^ and *FDR* = 4.9 × 10^−2^ for singlets and complexes, respectively). c Proportions of non-classical monocyte singlets and complexes in chronic asymptomatic and symptomatic individuals. Statistical significance was assessed using DCATS with a beta-binomial model (*P* = 1.6 × 10^−2^ for complexes). The same color legend was used for panel (a), (b) and (c). d-e UMAP projection and corresponding proportions of monocyte complex subtypes across individuals. The same color legend was used for panel (d) and (e). f Proportion of non-classical monocyte–T-cell complexes in chronic asymptomatic and symptomatic individuals. Statistical significance was assessed using DCATS with a beta-binomial model (*FDR* =2.0 × 10^−2^).

Fluorescence microscopy further confirmed the presence of several complex configurations, including monocyte–T cell–B cell assemblies and complexes in which lymphocytes were associated with a single monocyte (Supplementary Figure 2). Together, these observations provide independent visual evidence that the identified complexes represent bona fide immune cell complexes rather than technical artefacts.

### Flow cytometry validation of monocyte immune complex diversity

Based on the flow cytometry data generated to confirm the distribution of global cell types, we investigated initially filtered out multiplets to validate the diversity of immune complexes. Using a standardized antibody panel, we identified monocyte complexes in cryopreserved PBMCs samples (Fig. 6a). We were able to observe diverse monocytes complexes with T cell and B cell comprising all possible combinations of these three cell types (Fig. 6b, 6c). However, the antibody panel did not allow to capture platelets which may introduce bias in estimation of multiplet subtype proportions between the two technics. Overall monocyte complex proportion was higher in children with chronic infection than in symptomatic with a trend of association (P = 5.6 x 10^-2^). Classical monocytes (Fig. 6b) and non-classical monocyte multiplets (Fig. 6c) seem to be more frequent in children with chronic infection.

**Figure 6.**
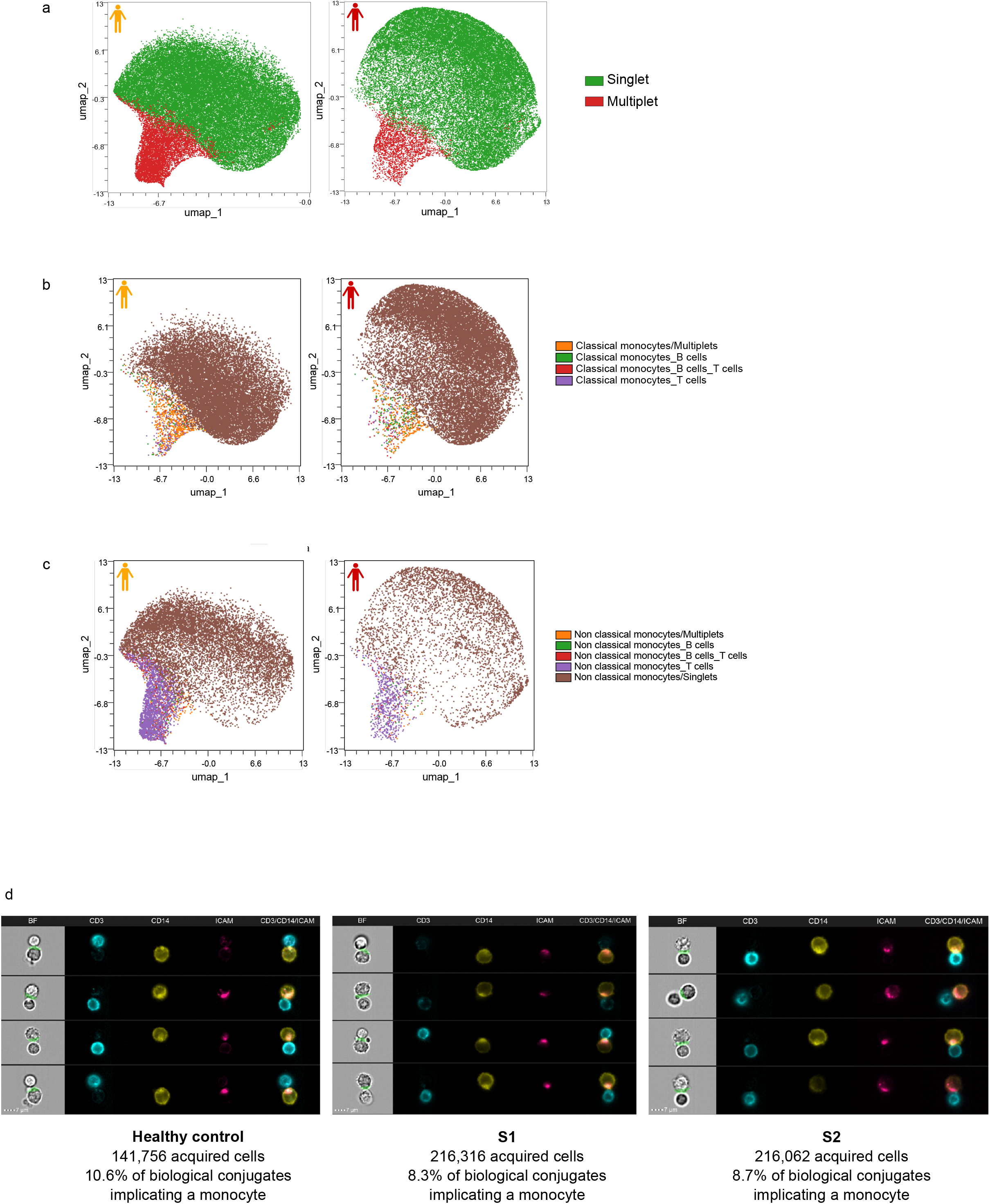
Unsupervised analysis with flow cytometry data obtained from PBMC of chronic asymptomatic and symptomatic patients was performed using OMIQ software on live monocytes. a UMAP plot of 13 000 monocytes from each of the 4 chronic (left) and the 3 symptomatic (right) patients reveal clusters of single cells (green) or multiple cells (red). b UMAP plot of 13 000 monocytes from each of the 4 chronic (left) and the 3 symptomatic (right) patients show clusters of classical monocytes in single cells (brown) or conjugates to B cells (green), T cells (purple), B and T cells (red). The total classical monocytes conjugated with other cells are shown in orange. c UMAP plot of 13 000 monocytes from each of the 4 chronic (left) and the 3 symptomatic (right) patients shows clusters of non-classical monocytes in single cells (orange) or conjugates to B cells (green), T cells (purple), B and T cells (red). The total non-classical monocytes conjugated with other cells are shown in orange. d Random galleries of CD3⁺ CD14⁺ conjugates with immune synapse identified by imaging flow cytometry. From left to right: bright field images with interface mask (in green), T cells [CD3-APC-Cy7], monocytes [CD14-PE], ICAM-1+ cells [ICAM-APC]. The final column shows CD3, CD14, ICAM-1 composite image overlays highlighting the polarization of ICAM-1 at the immune synapse interface.

### Imaging flow cytometry reveals physical interactions within monocyte–T cell complexes

Next, we investigated whether monocyte-T cell complexes displayed marker properties of physically interacting cells. Hence, we performed imaging flow cytometry which combines conventional flow cytometry with high resolution imaging, enabling direct visualization of cellular interactions. Classical monocyte-T cell complexes were analyzed in two symptomatic individuals previously included in the scRNA-seq experiment (S1 and S2) and in one healthy control. Cells were stained for CD3 to identify T cells, CD14 to identify classical monocytes, and ICAM-1 to discriminate between non biological and biological conjugates/complexes, using the gating strategy described in Supplementary Figure 3. Image analysis revealed classical monocyte–T cell complexes displaying a polarized accumulation of ICAM-1 at the cell-cell interface (Fig 6d), consistent with the formation and stabilization of monocyte-T cell immune synapses. These observations provide additional evidence that the monocyte-T cell complexes detected in peripheral blood reflect biologically meaningful cellular interactions. The health control, expected to exhibit a relatively resting immune state, exhibited 10.6% of monocyte-T cell complexes, compared with 8.3% and 8.7% in the two symptomatic individuals (Fig. 6d). These proportions were broadly consistent with those observed by scRNA-seq in the corresponding individuals. Given the limited number of samples analyzed, these observations do not allow statistical inference. Nevertheless, they provide preliminary evidence that the abundance and organization of monocyte–T cell interactions may vary according to clinical status.

Accordingly, rather than excluding these populations from downstream analyses, we considered them as candidate cellular interaction states and further investigated their potential contribution to the distinct immune landscapes associated with chronic asymptomatic and symptomatic malaria. These complexes may therefore represent candidate cellular signatures of immune responses associated with protection from symptomatic disease.

### Inflammatory transcriptional signatures distinguish symptomatic from chronic malaria

To identify transcriptional programs that discriminate between symptomatic and chronic malaria, we applied a machine learning framework to non-classical monocytes singlets and non-classical monocyte–T cell complexes (Fig. 7a). This approach, based on pseudo-bulk differential expression analysis, enabled the identification of dysregulated candidate genes and the establishment of gene signatures for the two cell populations (Fig. 7b, c). A minimal set of 20 genes using Logistic Regression achieved optimal classification performance for both non-classical monocytes and non-classical monocyte–T cell complexes (MCC = 1, F1-score = 1, and ROC-AUC = 1). Exhaustive permutation testing yielded an exact empirical P-value of 0.0286 for both cell populations, suggesting that the observed classification performance was unlikely to have occurred by chance, particularly for the complexes.

**Figure 7.**
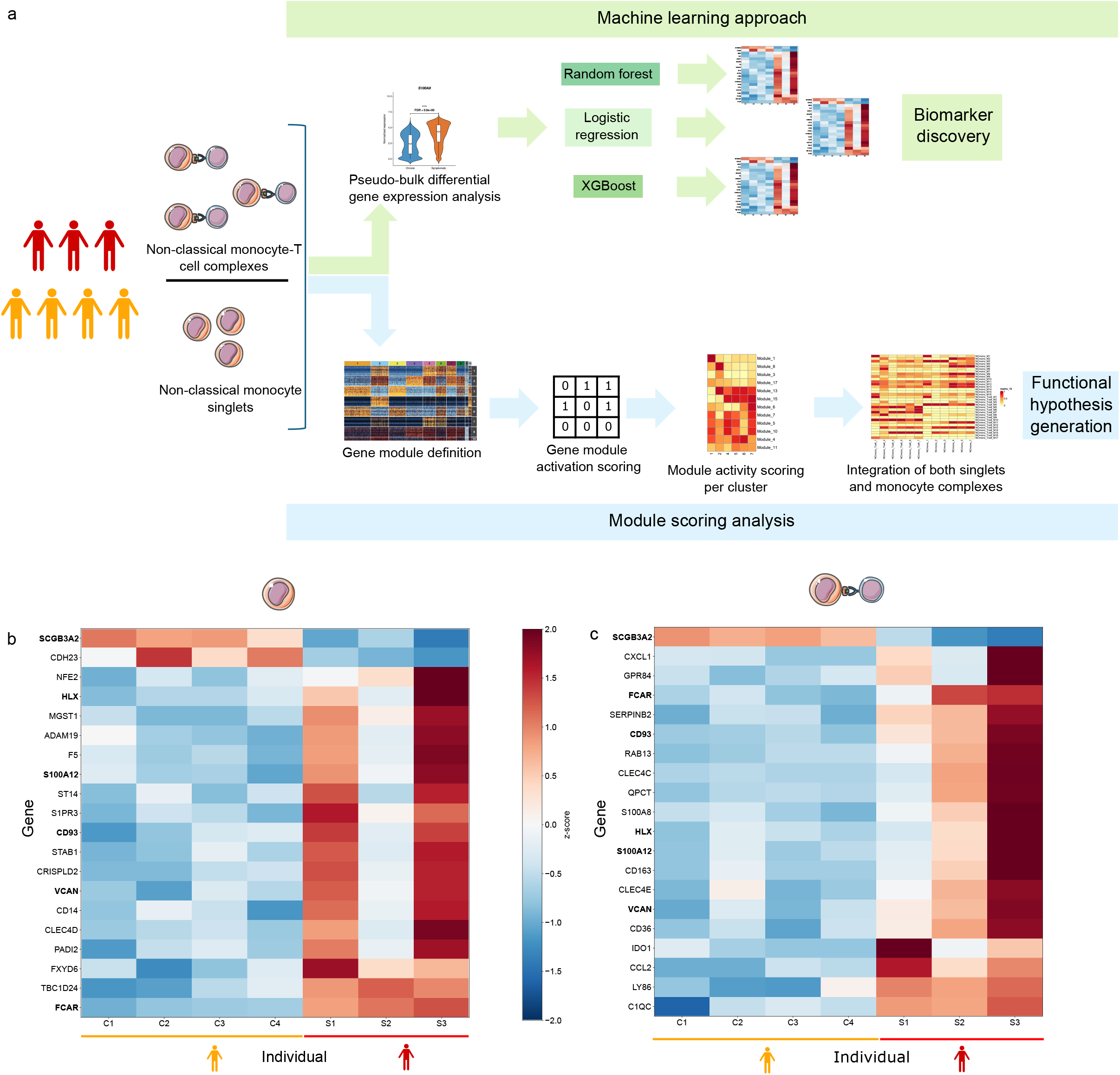
Machine learning-based identification of gene signature distinguished between the two clinical groups. a Schematic overview of the bioinformatic workflow used for signature identification using a machine learning approach and characterization of the transcriptional and population heterogeneity of non-classical monocytes and non-classical monocyte–T cells in chronic asymptomatic and symptomatic individuals. b-c Heatmap showing the top 20 genes identified by a machine learning approach based on logistic regression that discriminate non-classical monocytes. (b) and non-classical monocytes–T cell complexes (c) between chronic asymptomatic and symptomatic individuals. The same color scale is used for panels (b) and (c). Genes shared between (b) and (c) are shown in bold.

Most of the predictive genes contributing to the classification of symptomatic individuals were more highly expressed than in chronic individuals and were associated with inflammatory processes. This pattern indicates that symptomatic malaria is characterized by enhanced activation of inflammatory and innate immune pathways in both non-classical monocyte singlets and monocyte–T cell complexes. The comparison of the two top 20 gene signatures identified four shared genes: VCAN, S100A12, SCGB3A2 and FCAR that showed consistent directions of dysregulation across both cell populations (Fig. 7b and c). VCAN, S100A12 and FCAR, which are involved in inflammatory and innate immune responses, were upregulated in symptomatic individuals, whereas SCGB3A2, which has immunomodulatory and anti-inflammatory functions, was downregulated (Fig. 7b and c).

In addition, the heatmap revealed greater inter-individual heterogeneity among symptomatic individuals than among chronic individuals (Fig. 7b and c). Despite belonging to the same clinical category, symptomatic individuals displayed distinct transcriptional profiles, suggesting that different inflammatory states or trajectories may converge toward a broadly similar, yet heterogeneous symptomatic phenotype. In both monocyte singlets and monocyte–T cell complexes, we further observed a progressive gradient of gene expression across symptomatic individuals, consistent with variation in inflammatory intensity and potentially reflecting differences in clinical features such as fever or parasitemia. In contrast, chronic individuals exhibited more homogeneous transcriptional profiles. Together, these findings indicate that the biological heterogeneity of symptomatic malaria cannot be fully captured by differential expression analysis or by considering individual predictive genes alone.

### Contrasting inflammatory and immune-interaction states distinguish symptomatic and chronic malaria

We sought to identify subpopulations of either non-classical monocytes or monocyte-T cell complexes characterized by coordinated transcriptional programs that might be associated with a specific phenotypic group. Subsampling followed by gene module-based clustering identified clusters with varying proportions of cells from the two phenotypic groups (Fig. 8a and b). Despite inter-individual variability, overall, the non-classical monocytes from symptomatic individuals were overrepresented in clusters 4, 5, and 7, whereas cells from asymptomatic chronic individuals predominated in clusters 1, 2, and 3 (Fig. 8a). In contrast, no phenotype segregation could be observed among complexes (Fig. 8b).

**Figure 8.**
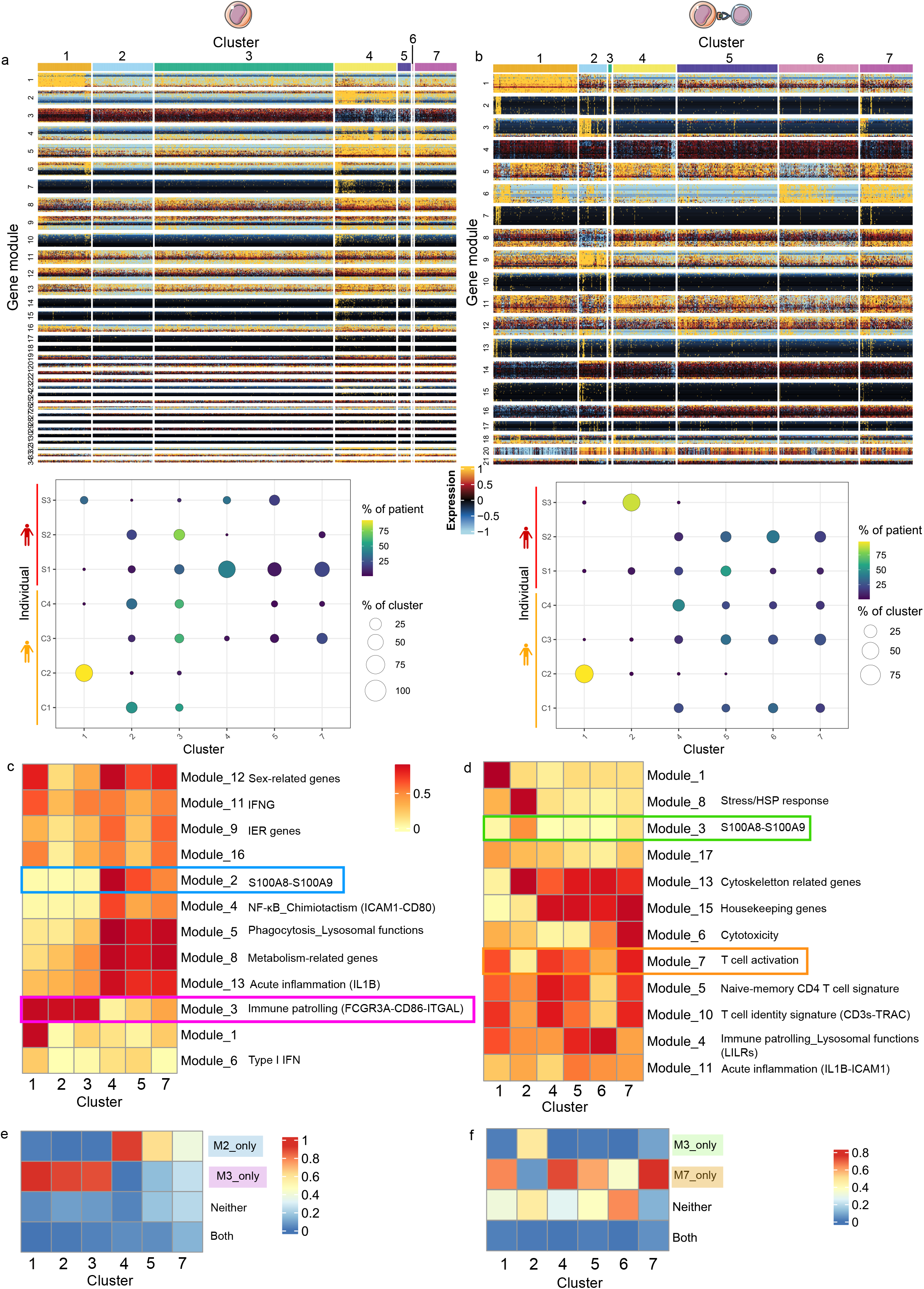
Inflammatory and surveillance transcriptional programs distinguish chronic asymptomatic and symptomatic individuals. a Heatmap showing modules of co-expressed genes used for unsupervised clustering of non-classical monocytes from chronic asymptomatic and symptomatic individuals (upper panel), and bubble plots showing the composition of the resulting clusters across individuals (lower panel). Dot size represents the proportion of cells belonging to each cluster, and color indicates the proportion of each individual to the corresponding cluster. b Heatmap showing modules of co-expressed genes used for unsupervised clustering of non-classical monocyte–T-cell complexes from chronic asymptomatic and symptomatic individuals (upper panel), and bubble plots showing the composition of the resulting clusters across individuals (lower panel). Dot size represents the proportion of cells in each cluster, and color indicates the proportion of each cluster within each individual. c, d Heatmaps showing module activity scores as the proportion of cells activating at least 50% of genes of a specific module across clusters identified in non-classical monocytes (c) and non-classical monocyte–T-cell complexes (d). IER, immediate early response genes. e, f Heatmaps showing the proportion of cells activating module 3 (M3), module 7 (M7) or both simultaneously in non-classical monocytes and (e) non-classical monocytes complexes (f). Modules enclosed in colored squares in panels (c) and (d) correspond to the same modules shown in panels (e) and (f), respectively. The same color scale is used for each pair of heatmaps: (a–b), (c–d), and (e–f).

Analysis of gene module activity scores further revealed distinct biological functions associated with specific clusters, particularly those related to cell-cell interaction, inflammation, cellular stress and T cell activation (Fig. 8c, 8d). Chronic individuals were characterized by higher activity of the FCGR3A, CD86 and ITGAL-associated program, suggesting a monocyte state with increased potential for immune surveillance and cell– cell interactions (Fig. 8c, Supplementary Table 7). In contrast, symptomatic individuals showed preferential activation of the S100A8, S100A9 and IL1B-associated inflammatory programs, together with increased activity of phagocytic, lysosomal and metabolism-related modules (Fig. 8c). These findings suggest that symptomatic malaria is associated with a broader inflammatory and functionally activated myeloid state, whereas chronic malaria is associated with a distinct monocyte state characterized by immune patrolling and cell–cell interaction programs. These contrasting activity profiles support the hypothesis of a transition between transcriptional states associated with cellular interactions and those associated with inflammation within the different groups. Furthermore, the concurrent activity of modules 2, 5, 8 and 13 suggests that the inflammatory state may coexist with phagocytic, lysosomal or metabolism-related programs, indicating a coordinated activation of multiple effector functions.

In non-classical-T cell complexes, sample S3, which developed symptomatic malaria with the most severe clinical manifestations, showed prominent activation of stress and inflammation modules, together with reduced T cell activation, despite the presence of interactions between non-classical monocytes and T cells (Fig. 8d). Overall, these observations suggest that inflammation may be associated with reduced cell-cell interaction in symptomatic individuals and that cellular stress and inflammation may also modulate or constrain T cell activation within monocyte–T cell complexes, particularly in severe symptomatic disease.

### Opposing inflammatory and cell-interaction programs at the cellular level

To assess whether inflammatory and surveillance-interaction programs were mutually exclusive, we evaluated the extent to which individual non-classical monocytes simultaneously activated modules 2 and 3 using a score reflecting the proportion of cells co-activating both modules (Fig. 8e). We showed that cells with activated module 2 did not activate module 3 and vice versa, indicating an inverse relationship between these transcriptional programs at the single-cell level. Notably, cells from symptomatic individuals activated the S100A8–S100A9-containing inflammatory module 2 but rarely activated the FCGR3A–CD86–ITGAL-associated module 3 concomitantly (Fig. 8e). We further confirmed the inverse relationship between inflammation (module 3) and T cell activation (module 7) at the single-cell level within non-classical monocyte-T cell complexes (Fig. 8f). Notably, at the individual level, cells from symptomatic children showed greater activation of inflammatory modules across clusters, regardless of cluster identity (Supplementary Figure 4). Taking together, these findings further support the existence of distinct and largely opposing transcriptional states, revealing their mutually exclusive activity at the cellular level. Cells from symptomatic malaria being primarily associated with an inflammatory state, potentially incompatible with cell–cell interaction establishment.

### The co-activation of *LILR* family genes and inflammatory programs could limit monocyte–T cell interactions in symptomatic individuals

Projecting the modules identified in non-classical monocyte singlets onto monocyte–T cell complexes revealed additional transcriptional programs that were not necessarily captured by gene expression correlations in the complex dataset but nevertheless showed coordinated activity. Among these, module 4 identified based on non-classical-T cell complexes, associated to immune surveillance and lysosomal functions also contain five *LILR* family genes (Fig. 9a). Module 4 was broadly activated in non-classical monocytes from both chronic and symptomatic individuals. Therefore, clusters 4, 5, and 7, which were predominantly composed of non-classical monocytes from symptomatic individuals, showed activation of S100A8-S100A9 inflammatory module 2 in addition to module 4 (Fig. 9a). This co-activation pattern was also observed at the single cell level: individual cells from clusters 4, 5 and 7 frequently showed simultaneous activity of both modules, whereas no cell from clusters 1, 2 and 3 which were predominantly composed of non-classical monocytes from chronic individuals displayed this combination of transcriptional programs (Fig. 9b). Consistently, no non-classical monocytes within complexes from either chronic or symptomatic children, showed concomitant activity of modules 2 and 4. Together, these observations suggest that the simultaneous activation of the *LILR* gene family and inflammatory programs define a distinct monocyte state that is not observed in monocyte–T cell complexes. This pattern further suggests that the functional state of the non-classical monocytes may influence its capacity to engage interactions with T cells.

**Figure 9.**
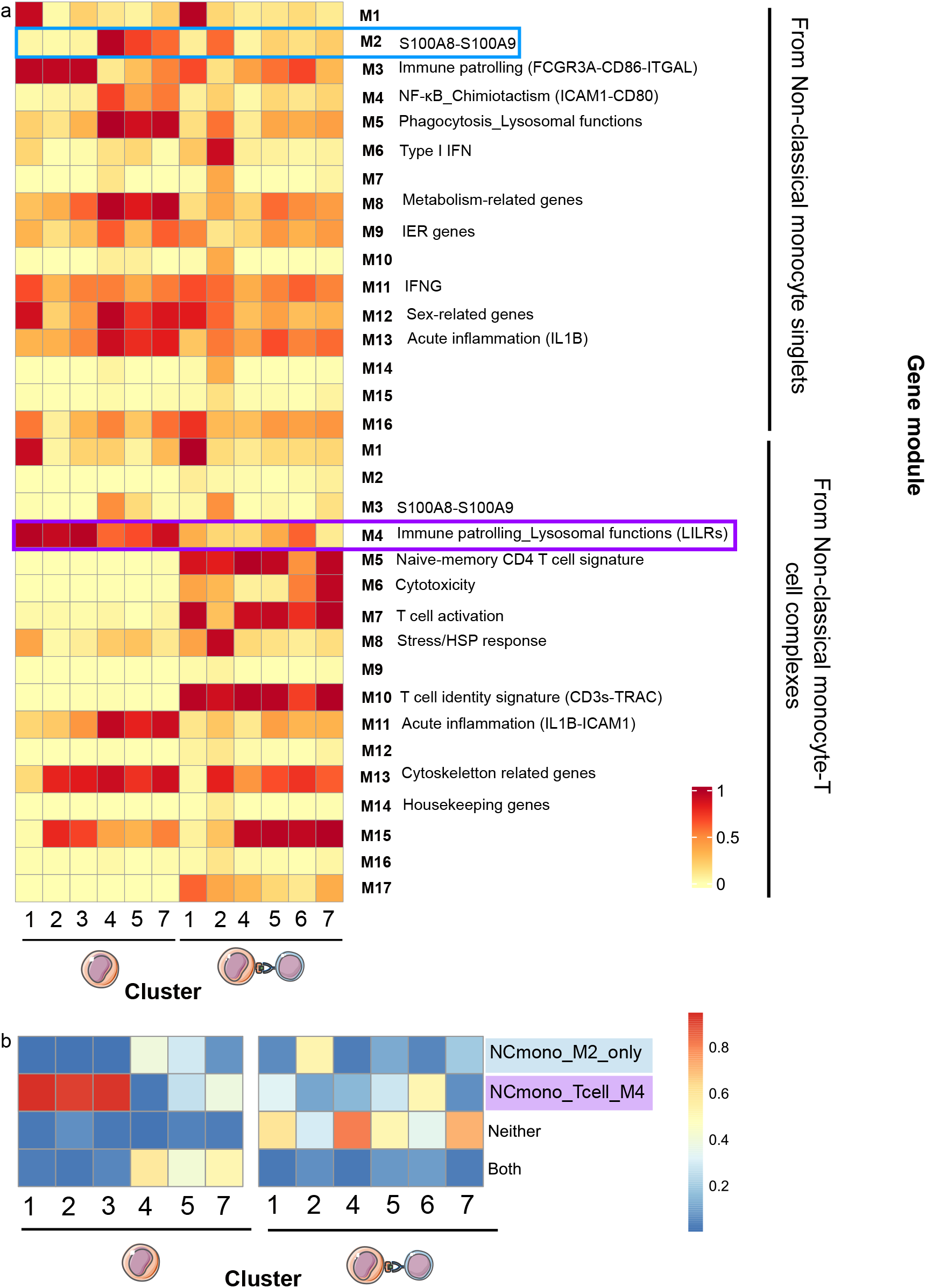
Co-activation of inflammatory signature and LILRs-containing programs in symptomatic individuals. a Heatmap showing the activity scores of gene modules identified in non-classical monocytes and non-classical monocyte–T-cell complexes across all clusters from both populations, enabling cross-population comparison of transcriptional programs. b Heatmaps showing the percentage of cells activating the S100A8-S100A9 module (M2), the immune patrolling lysosomal functions module (M4), neither of them or both simultaneously across clusters of non-classical monocytes and non-classical monocyte-T cell complexes.

## Discussion

In this study, we provide the first comprehensive characterization of the immune landscape of children with chronic and symptomatic malaria using single-cell RNA sequencing. Analysis of cellular composition revealed a higher fractional abundance of monocytes in symptomatic individuals, consistent with previous studies ^20,24^, highlighting monocytes as a potential contributor to the distinct immune profiles associated with clinical outcomes during malaria infection. The large number of cells analyzed per individual (mean average 9,933 cells, n = 7 individuals, total of 63,533 cells analyzed) and the bioinformatics strategy used enabled us to identify an unexpected subpopulation of cell complexes involving monocytes and T cells. Notably, monocyte-containing complexes comprising two or three cells were significantly more frequent in chronic individuals, despite the overall expansion of monocytes in symptomatic individuals. The main difference between chronic and symptomatic children was the frequency of interactions between non-classical monocytes and T cells. These findings suggest that monocyte abundance alone does not reflect their functional state and raise the possibility that physical interactions between monocytes-T cells and monocytes-T cells-B cells may be associated with the immune state characterizing chronic infection and limiting the development of clinical symptoms. Even if we observed a diversity of immune complexes by cytometry, we couldn’t confidently map all the cell types and conjugates due to the gating strategy used, highlighting the necessity to develop new approaches for the accurate identification of cellular interactions as proposed by Vonficht *et al.* ^25,26^. To our knowledge, this is the first scRNA-seq study that identifies cell populations corresponding to physical cell-cell interactions, whereas previous single-cell studies have primarily relied on computational approaches to infer putative cellular interactions ^20,22^. Importantly, our imaging analyses provided morphological evidence consistent with active monocyte -T cell interactions. We observed a polarized accumulation of ICAM-1 at the cell-cell interface, features consistent with the formation of an immunological synapse between monocytes and T cells ^27^. Interaction between antigen-presenting cells and antigen-specific T cells involves cytoskeletal reorganization and recruitment of adhesion molecules such as ICAM-1 and LFA-1, which contribute to the establishment and maintenance of antigen-specific immune responses ^28^. The existence and biological relevance of circulating T-cell-monocyte complexes have also been reported recently ^25,29^, and their frequency may vary in response to immune perturbations, including infection and immunization Polarized localization of LFA-1 and ICAM-1 at the cell–cell interface further support a potential role for these molecules in stabilizing T-cell–monocyte interactions ^26,30^. Consistent with our observations, a reduced frequency of T cell-monocyte complexes displaying synapse-like structures was reported in patients with active pulmonary tuberculosis compared with those with latent TB-infection and healthy controls, despite an increased frequency of CD14+ monocytes in active tuberculosis ^31^. Together, our findings support emerging evidence that T-cell–monocyte complexes may contribute to immune regulation during infection. In malaria, differences in the physical interactions between these cell populations may therefore represent an additional layer of immune regulation associated with clinical outcome.

This interaction phenotype appeared particularly related to the functional state of non-classical monocytes. These cells are commonly described as patrolling monocytes, with functions associated with vascular surveillance and the regulation or resolution of inflammatory responses ^32^. Their increased interactions with T cells in chronic individuals may therefore reflect a functional axis through which non-classical monocytes participate in the regulation of T-cell responses, potentially involving antigen presentation, adhesion, and immunoregulatory signaling pathways ^33^.To investigate whether additional molecular features of non-classical monocytes were associated with these distinct interaction patterns, we compared their transcriptional profiles both as individual cells and within monocyte–T-cell complexes. Pseudo-bulk analysis identified a 20-gene signature that discriminated symptomatic from chronic individuals in both individual non-classical monocytes and within complexes. Permutation analysis supports the observed separation, despite the small number of individuals. Overall, symptomatic individuals displayed increased expression of pro-inflammatory genes, including *S100A8*, *S100A12*, *FCAR*, and *VCAN*, consistent with previous transcriptomic studies of malaria ^18,20^. Interestingly, transcriptomic profiles were relatively homogeneous among chronic individuals, whereas greater inter-individual heterogeneity was observed among symptomatic children, possibly reflecting differences in clinical manifestations. Consistent with this interpretation, individual S3, who presented with the highest fever, displayed the strongest inflammatory transcriptional profile. Although these observations do not establish a direct relationship between monocyte transcriptional state and clinical severity, they suggest that the magnitude of monocyte activation may be associated with the degree of clinical inflammation.

At the single-cell level, we further identified an inverse relationship between inflammatory programs, characterized by *S100A8*, *S100A9*, and *IL1B*, and transcriptional features associated with patrolling and cell–cell interaction, including *FCG3A*, *CD8c* a co-stimulation molecule, and *ITGAL* (encoding LFA-1), an adhesion and signaling molecule both mandatory for antigen-presenting cell and T cell interactions. This pattern suggests that the inflammatory state observed in symptomatic individuals may be associated with a reduced capacity of non-classical monocytes to engage in T-cell interactions. Although non-classical monocyte–T-cell complexes were still detected in symptomatic individuals, their presence alone does not establish that these interactions are functionally equivalent to those observed in chronic individuals. One possibility is that the inflammatory and stress-associated states observed during symptomatic malaria alter the capacity of monocytes to establish or maintain efficient interactions with T cells. We therefore propose a working model in which the inflammatory state of monocytes influences their capacity to form and/or maintain functional interactions with T cells. In chronic individuals, the relatively lower inflammatory state may favor a patrolling phenotype and increase the capacity of non-classical monocytes to interact with T cells. This model provides a potential mechanistic link between monocyte inflammatory state, physical cell–cell interactions, and clinical phenotype, although the directionality and functional consequences of this relationship remain to be established.

In addition, the concomitant expression of inflammatory genes and members of the *LILR* gene family was predominantly observed in monocytes from symptomatic individuals Previous studies have shown that the calcium-binding proteins S100A8 and S100A9 can interact with LILRB1 ^34,35^. These observations raise the possibility that increased S100A8/S100A9–LILRB1 signaling may contribute to the altered functional state of monocytes in symptomatic malaria and, potentially, affect their interactions with T cells. Whether this pathway influences T-cell activation, antigen responsiveness, or the expression and function of HLA class I molecules remains to be determined. This observation is particularly relevant in the context of the immune-evasion mechanisms described for *P. falciparum*. A subset of RIFIN proteins expressed on the surface of infected erythrocytes can bind inhibitory LILR receptors, including LILRB1 and LILRB2. LILRB1-binding RIFINs have been shown to inhibit B-cell and NK-cell responses ^36^. LILRB2, which is expressed prominently by myeloid cells including monocytes, has also been identified as a RIFIN receptor ^37^. These findings raise the possibility that parasite-derived RIFIN–LILR interactions could contribute to the altered transcriptional state of monocytes during symptomatic infection and, potentially, influence their interactions with T cells. A study characterizing *rifin* expression patterns in these participants is currently in progress and will allow us to determine whether specific RIFIN repertoires are associated with the LILR-associated monocyte phenotype observed here.

Since monocyte-T cell-B cell complexes were almost absent from symptomatic individuals and their subtypes are unevenly distributed across individuals, they could not be further investigated. We can hypothesize that their rarity in symptomatic individuals could be a consequence of a low proportion of non-classical monocyte-T cell interactions, as such interactions enhance T cell-B cell interactions as demonstrated by Ansari et *al.* (2022) ^38^.

More broadly, our findings highlight the importance of considering physical cell–cell associations when interpreting single-cell datasets. Cell complexes captured during sample processing may contain biologically meaningful information that is lost when cells are analyzed exclusively as independent populations. Rather than representing technical artifacts alone, these complexes may provide a complementary view of the organization and regulation of immune responses during infection. In the context of malaria, differences in physical interactions between immune cell populations, in addition to changes in their abundance and transcriptional states, may therefore represent an additional layer of immune regulation associated with the development of clinical disease.

## Materials and methods

### Cohort and PBMC isolation

Children included in this study were recruited as part of a longitudinal cohort conducted in the Centre region of Cameroon. The design and clinical characteristics of the cohort are described in Fogang et al (manuscript in preparation). Briefly, children aged 5–15 years were treated with antimalarial and antiparasitic drugs at enrolment to clear pre-existing infections and were subsequently monitored weekly for *P. falciparum* infection by qPCR using finger prick blood samples. Venous blood samples were collected either at the onset of symptomatic malaria or during persistent asymptomatic infection 3 to 4 weeks after the first detection of *P. falciparum*. Peripheral blood mononuclear cells (PBMCs) were isolated by Lymphoprep density-gradient centrifugation, cryopreserved in liquid nitrogen, and processed using an identical protocol for all samples. For the present study, PBMCs from seven children (three symptomatic and four chronically asymptomatic infected) were selected for single-cell RNA sequencing and flow cytometry analyses. Demographic and clinical characteristics of these individuals are summarized in Supplementary Table 1.

### Ethics statement

All participants were enrolled following written informed consent from their parent or legal guardian. The study was approved by the ethical review committees in Cameroon (Comité national d’éthique de la recherche pour la santé humaine, N 2023/04/1534/CE/CNERSH/SP, 10 June 2021).

### Sample Fixation

Cryopreserved PBMC samples were thawed in RPMI pre-warmed to 37°C and supplemented with 15% FBS. Cells were centrifuged and resuspended in fresh medium to determine cell concentration and viability by Trypan blue exclusion using a Malassez counting chamber with an optical transmission microscope and a Countess Automated Cell Counter. Samples with a viability greater than 80% were subsequently fixed using the Chromium Next GEM Single Cell Fixed RNA sample Preparation kit (ref. 1000414) according to the manufacturer’s instructions (User Guide Fixation of Cells C Nuclei for Chromium Fixed RNA Profiling, CG000478, Rev C). Cells were visualized before and after fixation using an EVOS M5000 imaging system.

### 10x Genomics Chromium GEM library preparation and sequencing

Single-cell RNA sequencing was performed using the droplet-based 10x genomics platform using the chromium Fixed RNA kit, Human transcriptome (ref 1000475), Chromium Next GEM Chip Q Single cell Kit (ref 1000422) and Dual Index kit TS set A (ref 1000251) according to manufacturer’s instructions (CG000527, Rev E). Library preparation was carried out at the TGML platform in Marseille. Briefly, samples were multiplexed by four using probes containing cell barcodes. The number of cells loaded for each sample onto a 10x chip was determined according to the manufacturer’s recommendations (10x Genomics) to target a recovery of approximately 10,000 cells per sample. Cells were partitioned in a gel bead containing emulsion by the Chromium Controller. Libraries were sequenced using the Illumina NovaSeq 6000 sequencing platform.

### Single-cell RNA-Seq data processing

Following sequencing, BCL files were converted into FASTQ files using bcl-convert (v4.2.7) samples were demultiplexed using Cellranger multi (Cell Ranger v8.0.1, 10x Genomics) ^39^. Only cells expressing more than 200 features expressed in at least 3 cells were kept.

Analysis of the scRNA-Seq data was performed using the Seurat package (v4.3.0) and the standard R package (v4.2.2) ^40^. Each sample was analyzed separately at the beginning using the following standardized procedure. A step to flag cells was applied based on UMI count (maximum of 40 000 UMI per cell), feature count (maximum of 6500 features per cell) and percentage of mitochondrial genes expressed (maximum of 10%). Note that ribosomal gene expression was not assessed during this quality control step, as the 10x Flex probe panel does not include probes targeting these transcripts. The data was normalized using NormalizeData (LogNormalize). Gene expression values were centered (mean-centered per gene) but not variance-scaled using ScaleData. The 2000 most variable genes were then selected using FindVariableFeatures for principal component analysis (PCA). The first 30 principal components were then used for Uniform Manifold Approximation and Projection (UMAP) embeddings calculation and clustering based on Leiden algorithm from the FindlClusters function. For cluster identification, differential gene expression analysis across clusters was performed using FindAllMarkers (VST)with a Wilcoxon method. Clusters were then identified using both manual annotations based on literature and databases (PanglaoDB ^41^, Azimuth ^40^, ImmunexUT ^42^, CellMarker2.0 ^43^, SingleR ^44^, and previously reported immune cell signatures ^21,22^. A doublet identification tool, DoubletFinder ^45^ was then used to identify potential doublets and annotate them subsequently.

### Cell subtype refinement using SciGeneX

To subcluster and annotate rare cell populations, the SciGeneX (Single-cell informative Gene eXplorer) tool (v1.4.11) ^46^, was applied using the following standardized workflow. Each sample was analyzed separately within each major cell population (Naïve-memory B cell, monocyte, Effector-memory CD4 T cell, Effector-memory CD8 T and NK cells). Subsampling was performed based on the cell-type annotations obtained from the initial Seurat analysis. The single-cell RNA-seq analysis pipeline described in the previous section (Single-cell RNA-Seq analysis) was then applied on the resulting subset of cells to identify transcriptionally distinct subclusters. Next, SciGeneX tool was applied to identify modules of co-expressed genes within the subclustered populations. Only modules containing at least four genes were retained for downstream analysis. Genes present in these modules were subsequently used for data scaling, followed by reapplication of the standardized Seurat workflow from the FindVariableFeatures step. SciGeneX was then applied a second time to further refine the modules of co-expressed genes. Final cluster annotations were assigned based on the combination of gene modules characterizing each cluster and canonical cell-type markers identified among the co-expressed genes within these modules. Module scores for selected SciGeneX modules of interest were computed using the AddModuleScore tool from Seurat.

### Differential cell composition analysis using DCATS

Seurat v5.1.0 and R v4.4.0 were used for the analyses. To account for sampling variability and the dependence among cell-type proportions within each sample, differential cell composition was assessed using the DCATS tool ^47^, which implements a beta-binomial model for statistical significance testing. This approach was used to identify differences in the relative abundance of cell populations between the clinical groups.

### Global merging of monocytes and Harmony integration

Monocytes from all individuals were merged into a single dataset, and batch effects were corrected using Harmony. SciGeneX was then applied separately to the integrated non-classical monocyte and non-classical monocyte–T cell complexe datasets. The gene co-expression modules identified by SciGeneX were used to perform unsupervised clustering. Based on the resulting module expression heatmaps generated using the ComplexHeatmap package ^48^, seven clusters were defined independently for non-classical monocytes and non-classical monocyte–T cell complexes.

### Module activity scoring per cell

For this analysis, only modules containing at least 10 genes and exhibiting a minimum gene-expression standard deviation of 0.15, as determined in the previous step, were retained.

For each gene, the mean expression level was calculated across all cells within the corresponding cell population. Gene activity was then binarized at the single-cell level. For each cell, a gene was assigned a binary activity value of 1 if its expression level exceeded the mean expression of that gene across the cell population, and 0 otherwise.

For each module in each cell, a module activity score was calculated as the proportion of activated genes within the module (i.e. Genes with a value of 1).

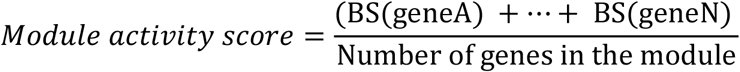

where BS denotes the binary gene activity score (0 or 1), and N is the total number of genes in the module.

A module was considered active in a given cell when its module activity score was greater than or equal to 0.5. Accordingly, active modules were assigned to a value of 1, whereas inactive modules were assigned toned a value of 0.

### Module activity scoring per cluster

For each cluster, module activity was defined as the proportion of cells within the cluster exhibiting activation of a given module.

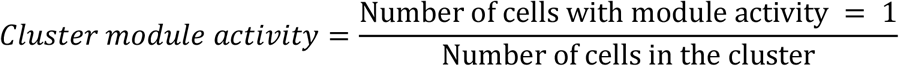

Thus, cluster-level module activity scores range from 0 to 1 and represent the fraction of cells activating a given module.

Modules were identified based on Gene-Ontology (GO) over-representation analysis using gProfiler, Reactome pathway enrichment using enrichR (Reactome 2026), and a review of the relevant literature.

### Pseudobulk Generation and Gene Ranking

Single-cell RNA sequencing data were aggregated into patient-level pseudobulk expression profiles by summing raw gene counts for each patient within each annotated cell type. This approach preserves the patient as the biological unit of replication while reducing cell-to-cell variability and accounting for the non-independence of cells from the same patient. Pseudobulk samples containing fewer than 20 cells were excluded from downstream analyses to ensure adequate cellular representation. In addition, genes with fewer than 10 total counts across all pseudobulk samples were removed prior to differential expression analysis to minimize the influence of lowly expressed features.

Differential expression analysis was performed independently for each cell type using the DESeq2 framework, which fits a negative binomial generalized linear model, with disease status as the explanatory variable ^49^. To prioritize candidate biomarkers for subsequent machine learning analyses, genes were ranked using a composite score integrating both statistical significance and the magnitude of differential expression as measured by the absolute log2 fold change.

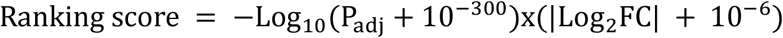

This ranking was used exclusively for feature selection and was not intended to guide the biological interpretation of differential expression results.

### Machine Learning Classification

Normalized pseudobulk expression matrices, transformed into log-transformed counts per million (logCPM), were used as input for supervised machine learning analyses. To evaluate the impact of feature dimensionality on predictive performance, classifiers were trained using progressively reduced feature subsets ranging using the top 40, 30, 20, 10 and 5 ranked genes.

Three complementary supervised learning algorithms were evaluated: Random Forest ^50^, Logistic Regression ^51^, and Extreme Gradient Boosting (XGBoost)^52^.

Model generalizability was assessed using Leave-One-Out Cross-Validation (LOOCV), in which a single patient was iteratively designated as the independent test sample while the remaining patients were used for model training. This strategy ensured that the predictive performance reflected the model’s ability to generalize to individuals who had not yet been observed.

Classification performance was evaluated using multiple complementary metrics, including accuracy, precision, recall, specificity, F1-score, Matthews Correlation Coefficient (MCC), and the area under the receiver operating characteristic curve (ROC-AUC). Given the limited sample size, MCC was selected as the primary metric for model evaluation because of its robustness to class imbalance and small datasets ^53^. Models were ranked according to MCC, followed by ROC-AUC and F1 score, and the three highest-ranked configurations were retained for downstream analyses.

### Statistical Validation and Model Interpretation

To determine whether the observed predictive performance exceeded that expected by chance, statistical significance was assessed using permutation testing. Given the limited number of individuals (n = 7; 4 Chronic and 3 Symptomatic), the 200 random permutations were replaced by an exhaustive patient-level permutation approach. All 35 possible group assignments preserving the observed class sizes (*^7^*) = *35* were evaluated using the same cross-validation procedure. Exact empirical P-values were calculated as the proportion of configurations achieving predictive performance equal to or greater than that observed with the original clinical labels.

The biological relevance of the selected models was subsequently investigated using SHapley Additive exPlanations (SHAP), which quantify the contribution of individual genes to model predictions ^54^. SHAP values were used to rank genes according to their contribution to the predictive models, with higher absolute SHAP values indicating a greater influence on model predictions. Gene level importance scores were finally integrated with differential expression statistics to facilitate the identification and biological interpretation of candidate disease-associated biomarkers.

### Theoretical calculation of multiplet proportions

The expected proportion of doublets and triplets from the 10x Genomics were estimated from the multiplet rate reported by 10x Genomics using a Poisson model. Based on the observed data, the expected numbers and proportions of neotypic and embedded multiplets were subsequently estimated using a binomial model.

### Fluorescence microscopy

Fixed cells were centrifuged at 850 g for 5 minutes and resuspended in PBS 1X at approximately 1 million of cells by ml. Antibodies (anti-human CD3-FITC, CD14-PE, CD16-vioblue and CD19-APC) were added and incubated for 20 minutes on ice and then washed twice with staining buffer (PBS containing 0.5% FBS and 2 mM EDTA pH8.0). The cells were observed with an inverted Zeiss Z1 automated microscope (Carl Zeiss).

### Flow cytometric cell phenotyping

PBMCs from the seven individuals were thawed in a pre-warmed complete medium (RPMI + 10% FBS **+** 1mM Sodium Pyruvate + 2mM L-glutamine) at 37°C. Dead cells were identified using a fixable viability dye (FVS 440 UV, BD Biosciences) and Fc receptors were blocked using human serum. Cells were then stained with chemokine receptor antibodies for 1 hour at room temperature. Following washing steps, surface staining was performed for 30 min at 4°C. Finally, cells were fixed using a 4 % paraformaldehyde solution (Electron Microscopy Sciences). Flow cytometry data were analyzed using FlowJo software (BD Biosciences) for conventional analyses and OMIQ for unsupervised clustering and dimensionality reduction.

### Imaging flow cytometry

PBMCs from the two symptomatic donors (S1 and S2) and one healthy donor (obtained from French Blood Establishment, Etablissement Français du Sang) as a control, were thawed in a pre-warmed complete medium at 37°C. Surface staining was conducted for 30 minutes at 4°C, followed by two washing steps. Cells were then resuspended in phosphate-buffered saline (PBS) supplemented with 2% fetal bovine serum (FBS). Immediately after filtration, samples were acquired on ImageStreamX MarkII (ISX, Amnis-EMD Millipore) at 40X magnification. Acquisition was performed at low speed to optimize image sensitivity. Single-stained samples were collected under the same settings and used as compensation controls to generate a compensation matrix. For each sample, a minimum of 200.000 events were acquired ensuring a sufficient number of biological CD3+ CD14+ conjugates.

Image analysis was performed using the ISX Data Exploration and Analysis Software (IDEAS, version 6.6.64.0, EMD Millipore). Multiplet cells in focus were discriminated from single cells upon the aspect ratio versus area plot of the bright-field channel. Morphology masks were generated based on CD3+ and CD14+ staining, and CD3+ CD14+ cells were identified based on the fluorescence intensity within the corresponding masks.

Only conjugates containing a single CD3⁺ cell and a single CD14⁺ cell were selected. An “interface” mask was created to determine the area of interaction between the CD3⁺ T cells and the CD14⁺ cells. The ‘delta centroid XY’ parameter between the interface mask and the CD3⁺ mask was used to select CD3⁺ CD14⁺ cell pairs with a defined contact zone, ensuring that the identified conjugates had effectively interacted. The intensity of

ICAM-1 was calculated within the interface mask and on the CD14⁺ cell. To normalize the data, the ratio of ICAM-1 intensity at the interface to that on the overall CD14⁺ cell was calculated. A high interface ratio indicating the recruitment or the enrichment of ICAM-1 at the CD3⁺ CD14⁺ interface, was therefore used to identify biologically relevant CD3⁺ CD14⁺ conjugates.

### Antibodies used in this study

CD45RA-BV480 (clone HI100), CD28-BV570 (clone CD28.2), CD138-BV605 (Clone MI15), CD56-BV650 (clone NCAM16.2), CCR6-BV711 (Clone 11A9), CRTH2-BV750 (Clone BM16), CCR7-BV785 (Clone G043H7), CD14-BV421 (Clone TÜK4) or CD14-PE (Clone 63D3.rMAb) (ISX), PD1-BB630, CXCR5-BB660 (clone RF8B2), IgM-PercP Cy5.5 (clone MHM-88), CD57-BB790 (clone NK-1), CD36-FITC (clone REA760), CD25-PE CF594 (clone BC96), CXCR3-PE Cy5 (Clone 1C6/CXCR3), CD19-PE Cy5.5 (clone SJ25-C1), TLR2-PE (clone REA109), CXCR6-PE Cy7 (clone K041E5), CD38-BUV395 (clone HIT2), CD4-BUV496 (clone SK3), CD8-BUV563 (clone RPA-T8), IgD-BUV615 (clone IA6-2), CD127-BUV661 (clone HIL-7R-M21), CD27-BUV737 (clone M-T271), CD69-BUV805 (clone FN50), CD3-APC Cy7 (Clone REA613 or UCHT1 (ISX)), HLA-DR-AF700 (clone L243), CD16-APC (clone 3G8), ICAM-APC (Clone HCD54)(ISX), CD3-FITC (clone REA613), CD14-PE (clone REA599), CD16-vioblue (clone REA423), CD19-APC (clone REA675).

## Data availability

All data generated or analyzed during this study are included in this article and its Supplementary Information files or are available from the corresponding author upon reasonable request.

The raw sequencing data and processed Cell Ranger outputs generated in this study have been deposited in the NCBI Gene Expression Omnibus (GEO) under accession number GSE348164: https://www.ncbi.nlm.nih.gov/geo/query/acc. The dataset includes samples GSM10067368 (PBMC_chronic_C2), GSM10067369 (PBMC_symptomatic_S3), GSM10067370 (PBMC_symptomatic_S2), and GSM10067371 (PBMC_chronic_C1_C3_C4_symptomatic_S1).

## Code availability

Associated analysis scripts are available at the GitHub repository release v1.0: MADBIO_Project v1.0. All analyses were performed in a containerized environment, and the corresponding Dockerfile is available in the same GitHub repository. All other data generated in this study are provided in the Supplementary Information. Source data are provided with this paper.

## Acknowledgements

This work was supported by the French Agency for Research (Agence Nationale de la Recherche [ANR]) ANR-22-CE15-0025-01 ‘‘MADBIO’’ (to S.M.), ANR 18-CE15-0009-01 (to A.C.), International research projects (IRP) “MalariaOmics” (to S.M) and by the Institut National de la Santé et de la Recherche Médicale (Inserm). L.Q. and M.A. were supported by a PhD fellowship from the French Ministry of Higher Education and Research (MESR). The scRNA sequencing was performed by the TGML platform (Marseille, France), supported by grants from INSERM, GIS IBiSA, Aix-Marseille Université, and France Génomique. We thank Béatrice Loriod and Denis Puthier managers of the TGML platform. We thank Laurine Gil and Pierre Milpied from Centre of Immunology Marseille-Luminy (CIML) for access to their platform. We thank Pascal Rihet and Julie Bavais from TAGC and Olivier Theodoly from Adhesion and Inflammation lab (LAI), for their helpful discussions. We thank Romain Miranda-Capet and Anaïs Almayrac for technical assistance on flow cytometry experiments. We thank all individuals who participated in the study cohort. The authors also thank Aurélie Bergon from TAGC for help with data management and Louis Souville for assistance with data deposition. Some elements of the figures were adapted from Servier Medical Art (https://smart.servier.com), licensed under CC BY 4.0 (https://creativecommons.org/licenses/by/4.0/).

## Author contributions

S.M. conceived and supervised the project. A.C. and S.M. designed cohort recruitment and supervised sample management. A.C. provided financial support for sample recruitment. M.A., C.G., M.T. and S.M. performed cell fixation and scRNA-seq experiments; L.Q., M.A., A.A. and L.S., performed the bioinformatic analyses, L.Q., L.S., and S.M., carried out interpretation and conceptualization of data. B.T., and S.N., conducted and supervised clinical studies and sampling; C.C., and B.F., contributed to study design and sample management; S.Mé, V.D.E., and A.L.I., performed cytometry and imaging flow cytometry experiments. A.B., and N.B., supervised cytometry and imaging flow cytometry experiments. M.B.P., performed fluorescence microscopy experiments. L.Q., S.Mé., and S.M., prepared the figures. L.Q., and S.M., wrote the manuscript with input from all authors.

