## Supplementary figures and images for "Single-cell characterization of the human immune landscape reveals dysfunction in monocyte-mediated cellular interactions in symptomatic malaria"

### Supplementary Figure 1

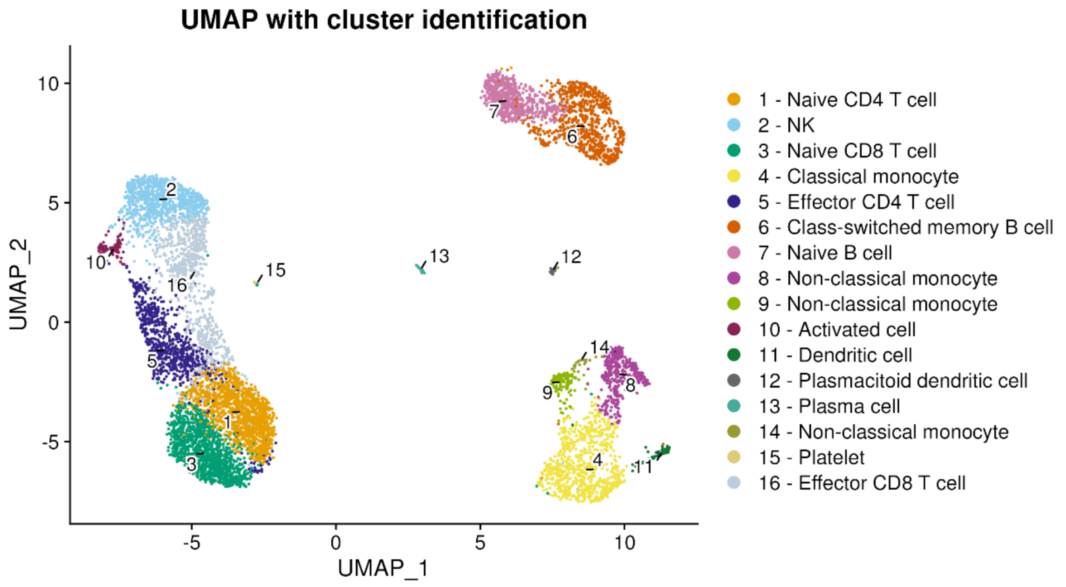

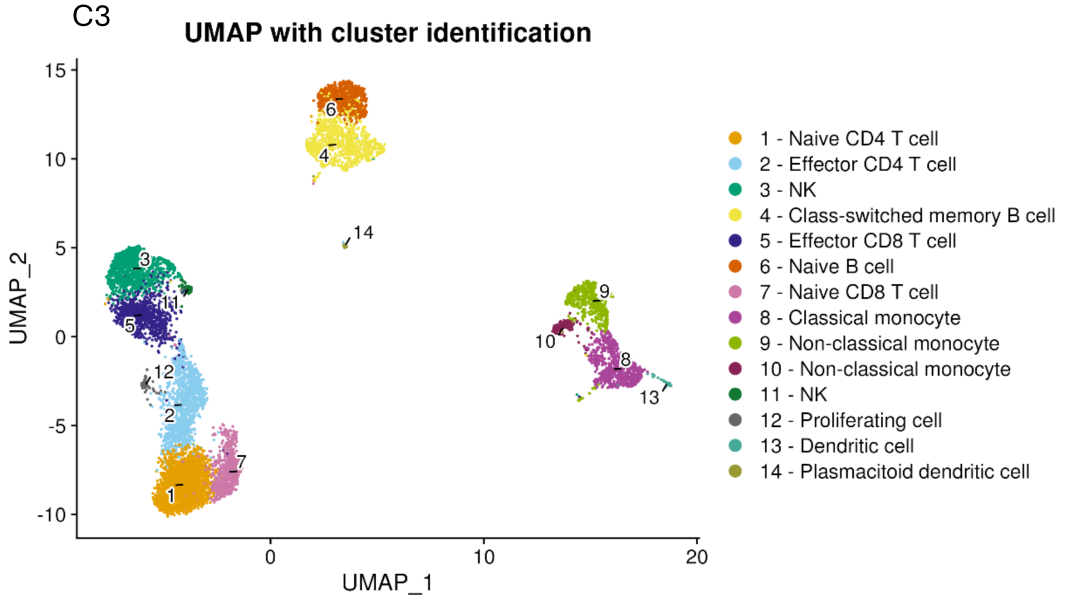

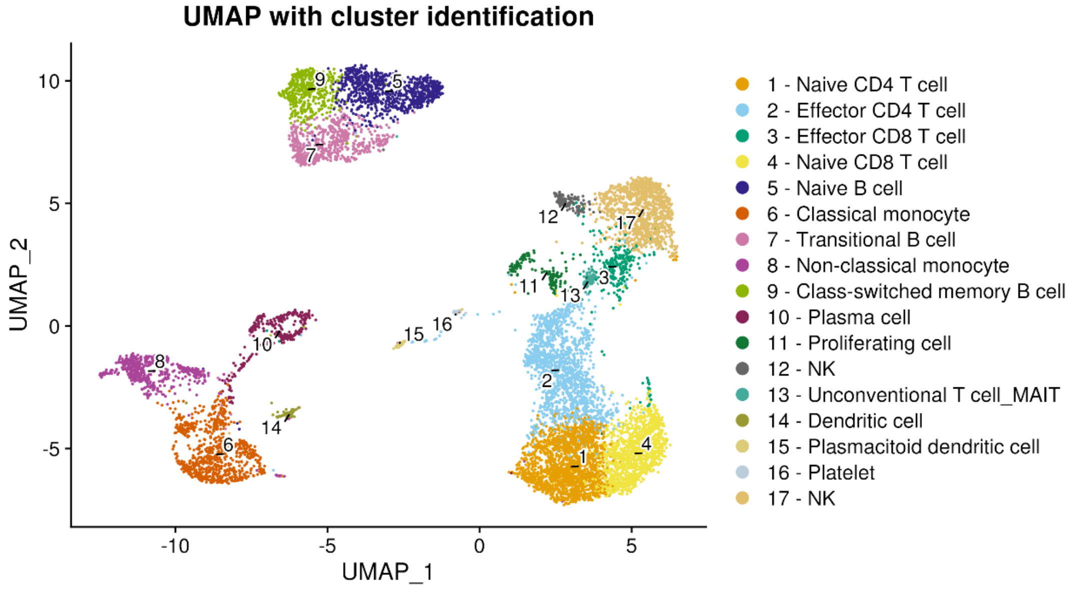

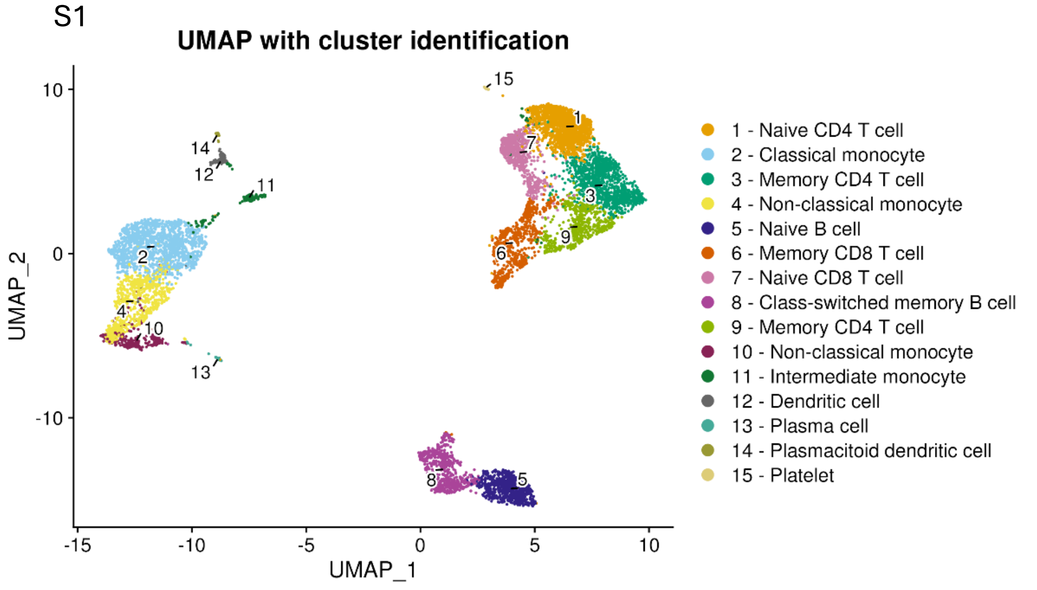

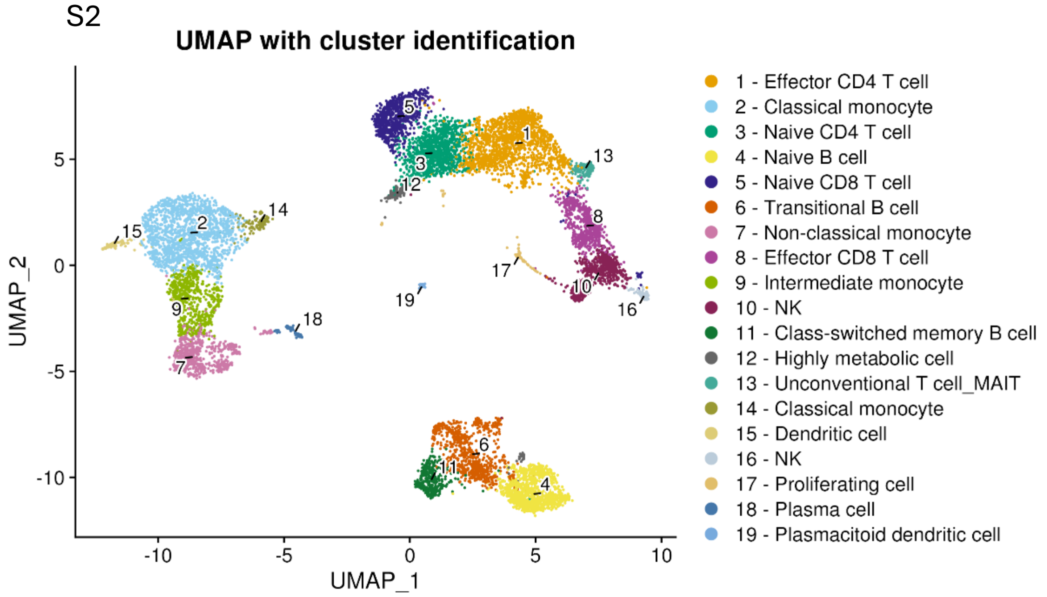

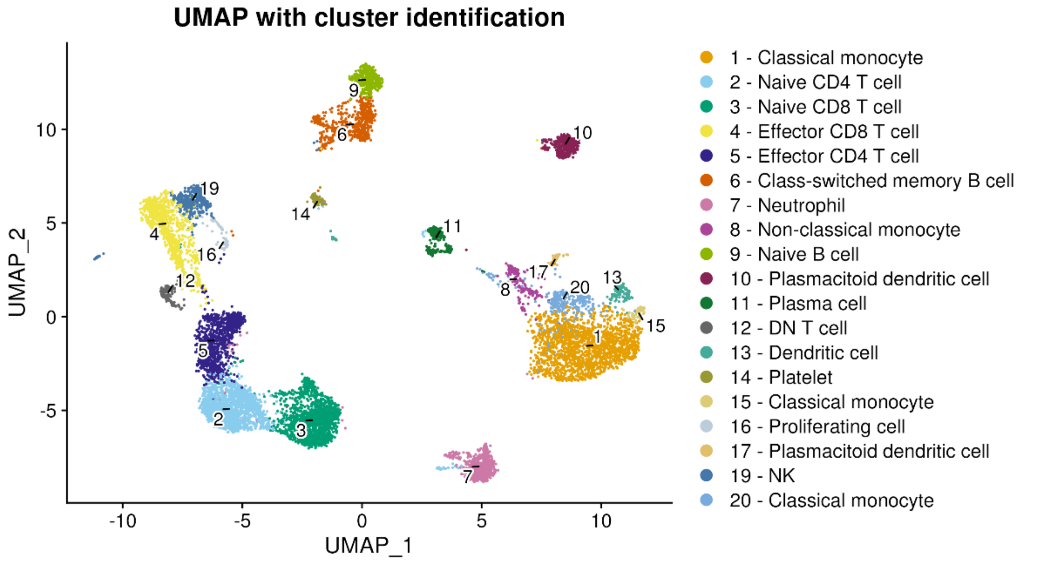

### Supplementary Figure 2

a

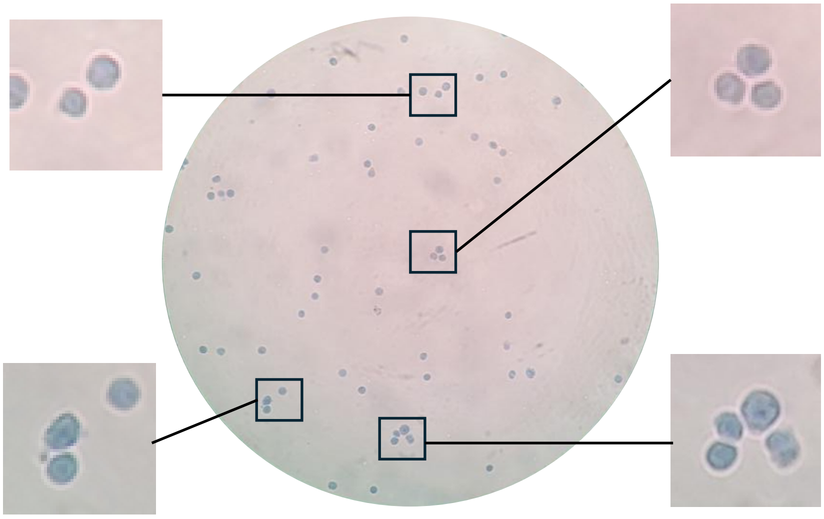

b

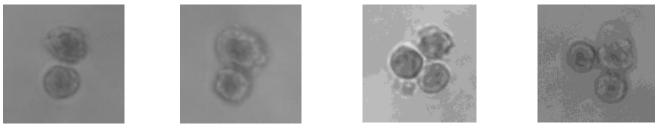

c

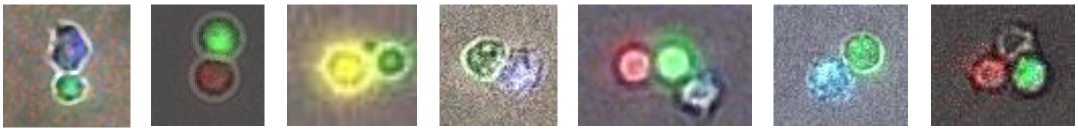

### Supplementary Figure 3

#### Gating strategy used for flow cytometry

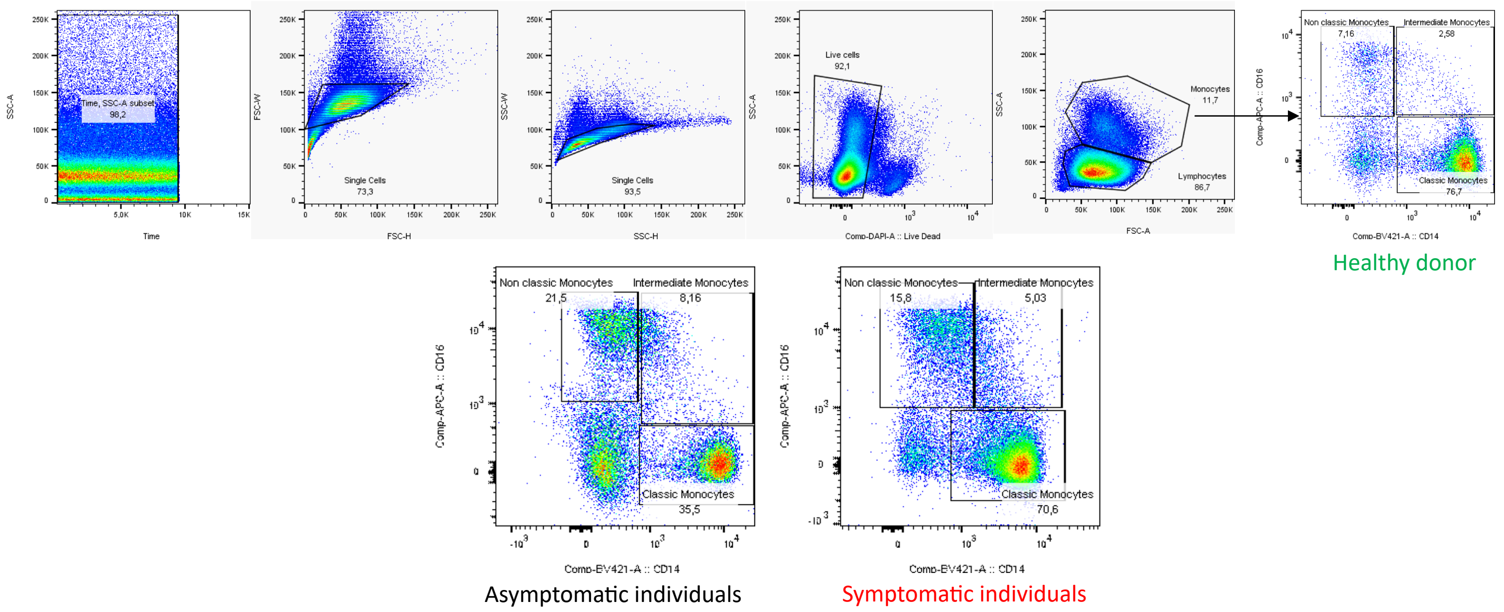


#### Gating strategy used for imaging flow cytometry

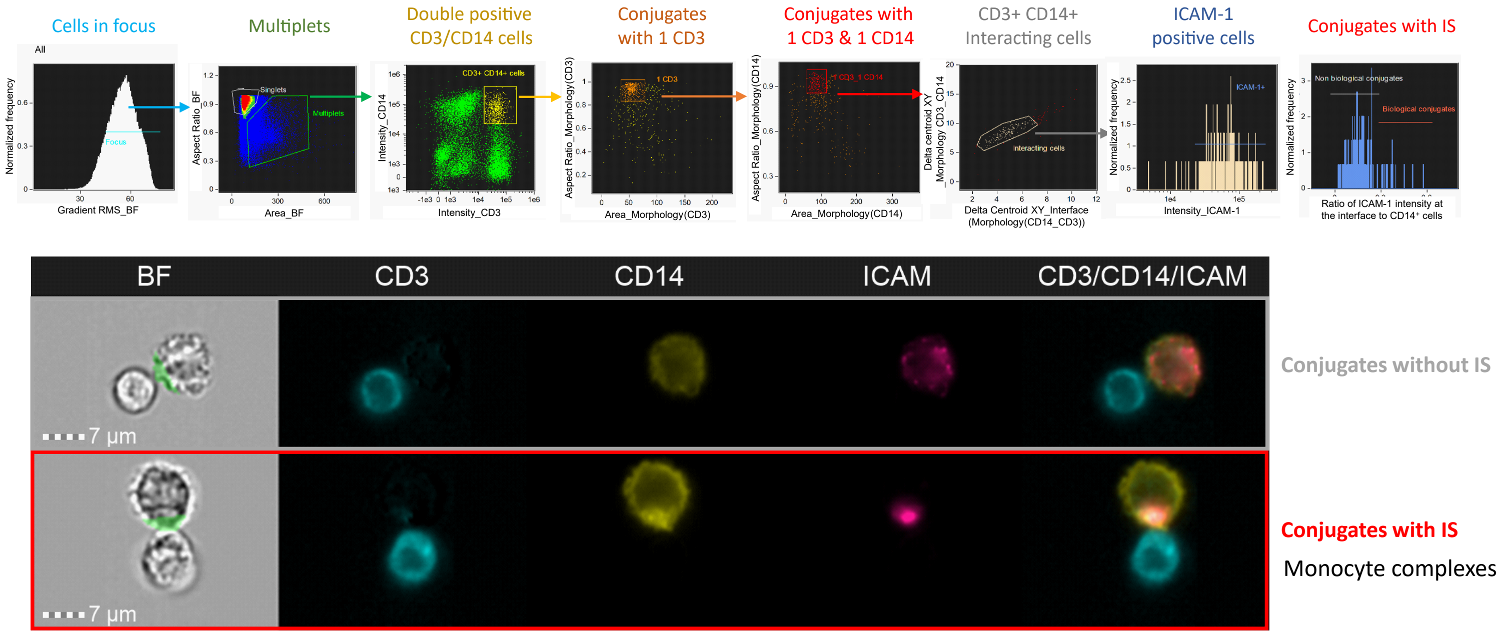

### Supplementary Figure 4

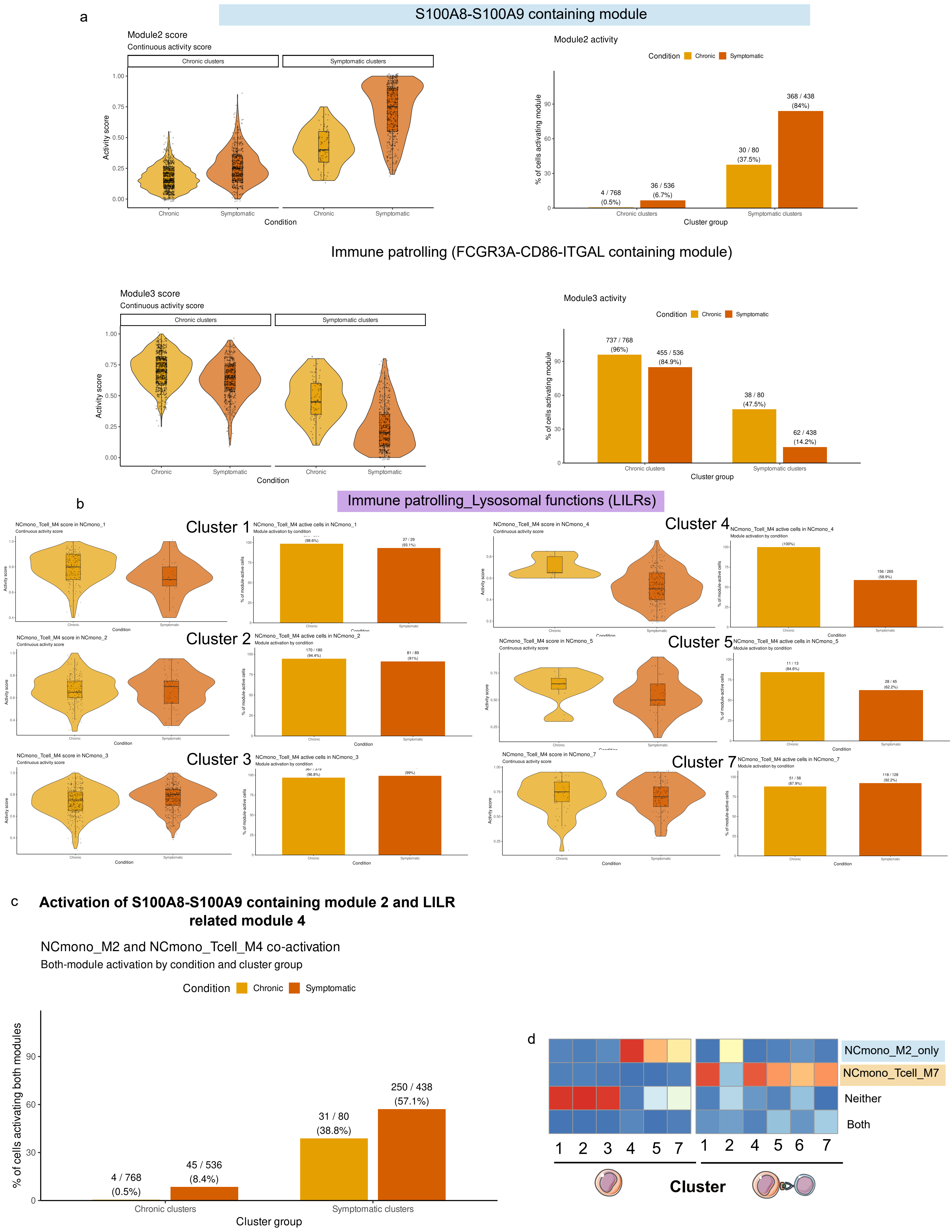
